# Comparisons of cell proliferation and cell death across life histories in the hemichordate *Schizocardium californicum*

**DOI:** 10.1101/2022.02.16.480686

**Authors:** Paul Bump, Margarita Khariton, Clover Stubbert, Nicole E. Moyen, Jia Yan, Bo Wang, Christopher J. Lowe

## Abstract

**Background:** There are a wide range of developmental strategies in animal phyla, but most insights into adult body plan formation come from direct-developing species. For indirect-developing species, there are distinct larval and adult body plans that are linked together by metamorphosis. Some outstanding questions in indirect-developing organisms include the extent to which larval tissue undergoes cell death during the process of metamorphosis and when the tissue that will give rise to the adult originates. Here we present patterns of cell proliferation and cell death during larval development, metamorphosis, and adult body plan formation, in the hemichordate *Schizocardium californium* to answer these questions.

**Results:** We identified distinct patterns of cell proliferation between larval and adult body plan formation of *S. californicum*. We found that some adult tissues proliferate prior to the start of any morphological metamorphosis. Additionally, we describe a genetic signature of proliferative cells with an irradiation approach that revealed markers shared between the life history states and others that are unique to larvae or juveniles. Finally, we observed that cell death is minimal in larval stages but begins with the onset of metamorphosis.

**Conclusions:** Cell proliferation during the development of *S. californicum* has distinct patterns in the formation of larval and adult body plans. However, cell death is very limited in larvae and begins during the onset of metamorphosis and into early juvenile development in specific domains. The populations of cells that proliferate and give rise to the larva and juvenile have a genetic signature that is more suggestive of a heterogeneous pool of proliferative progenitors versus a population of pluripotent set-aside cells. Taken together, we propose that *S. californicum* has a transformative metamorphosis that may be more representative of the development strategies that characterize metamorphosis in many metazoan animals.

## Background

The study of developmental biology has largely been informed by research in a few key model species that form their adult body plan during embryogenesis; a strategy termed direct development. However, this type of development is not representative of many animal groups where embryogenesis gives rise to a larva with a body plan distinct from that of the adult, a strategy which is called indirect development (1–3). In direct development, the adult is formed directly from the embryo, while in indirect development, embryonic processes give rise to a distinct larval body plan that later transforms into the adult. This transformation between larvae and adults is metamorphosis, a developmental event, with the loss of larval-specific structures and the emergence of adult structures, and is clearly mechanistically distinct between groups (4–10). The prevalence of this developmental strategy across bilaterian groups clearly demonstrates that a better mechanistic understanding of indirect development is critical for a fuller understanding of the developmental basis of body plan evolution.

The ocean provides a vast diversity of indirect developing strategies as many marine organisms first develop as larvae that will feed and grow before reaching metamorphosis (11, 12). In species such as gastropods with veliger larvae or polychaetes with trochophore larvae, the morphological difference between larval and adult body plans is not extreme, and metamorphosis represents a major shift in ecological niche but not a large morphological change (13–16). At the other end of the spectrum, as is found in some echinoids, larval and adult body plans can be radically different in organization with a “catastrophic metamorphosis” where the adult develops as a rudiment within the larva, and metamorphosis results in a complete reorganization of the body around new developmental axes in addition to the loss of larval structures (4,17,18). Similarly in some nemertean worms, with pilidium larva, the adult develops from several rudiments and metamorphosis is dramatic with the adult juvenile consuming the larval tissues (18, 19). However, metamorphosis in species with distinct larval and adult body plans does not always involve a segregated rudiment and cataclysmic metamorphosis — instead larval tissue seems to be remodeled rapidly into the adult without obvious drastic histolysis of the larval body plan (1,15,20). In this type of metamorphosis, the fate of larval tissue and origin of the adult body plan remains poorly understood. Indirect-developing hemichordates represent this type of metamorphosis and provide an opportunity to explore this type of developmental strategy.

Hemichordates are a phylum of worms composed of two classes, the solitary enteropneust worms and the largely colonial, tube-dwelling pterobranchs (4,21–24). While the position of hemichordates as sister to the echinoderms with a close relationship with chordates has been well established (25–28), new literature has challenged this position (29). Within the enteropneusts, one family, the Harrimaniiidae, are direct developers, while the other families, such as the Spengelidae and Ptychoderidae, are indirect developers with a distinct larval body plan called the tornaria. While morphological studies of tornaria larvae and their counterpart adult bodies can be traced back to the late 1800s (30), more recent morphological descriptions of larval and adult body plans have been carried out in a range of enteropnuest species; *Ptychodera flava* (31), *Balanoglossus misakiensis* (32), *Balanoglossus simodensis* (33) and *S. californicum* (34). In these species, the tornaria larva is formed directly following embryogenesis, while the benthic adult body plan forms by metamorphosis following an extended planktonic period (22, 28). These studies of hemichordate complex life cycles have largely been based on morphological characters, with some descriptive patterning studies (35–51). However, the cellular and developmental mechanisms driving the transformation of the larval body plan into the adult body plan remain largely uncharacterized. We do not know whether the adult is formed by transformation of larval tissues via differentiation and specification or by proliferation following large-scale larval tissue death.

To begin to address these questions, we characterized proliferation and cell death through the development and metamorphosis of *S. californicum* (Figure 1). We found distinct patterns of proliferation between larval and adult body plans. To then determine if there were distinct genetic markers of proliferative cells, and if those markers differed between life history stages, we also deployed an irradiation strategy to deplete proliferative cells and found a number of differentially expressed transcripts. When we considered the balance between cell proliferation and cell death, we found that cell proliferation of the adult body plan starts **prior** to metamorphosis. However, the start of a clear morphological metamorphosis corresponded with an increase in cell death. Along with the timing of these processes, the patterns of cell proliferation and cell death provided insights into the cellular contribution and transformation that occurs across metamorphosis.

**Figure 1.**
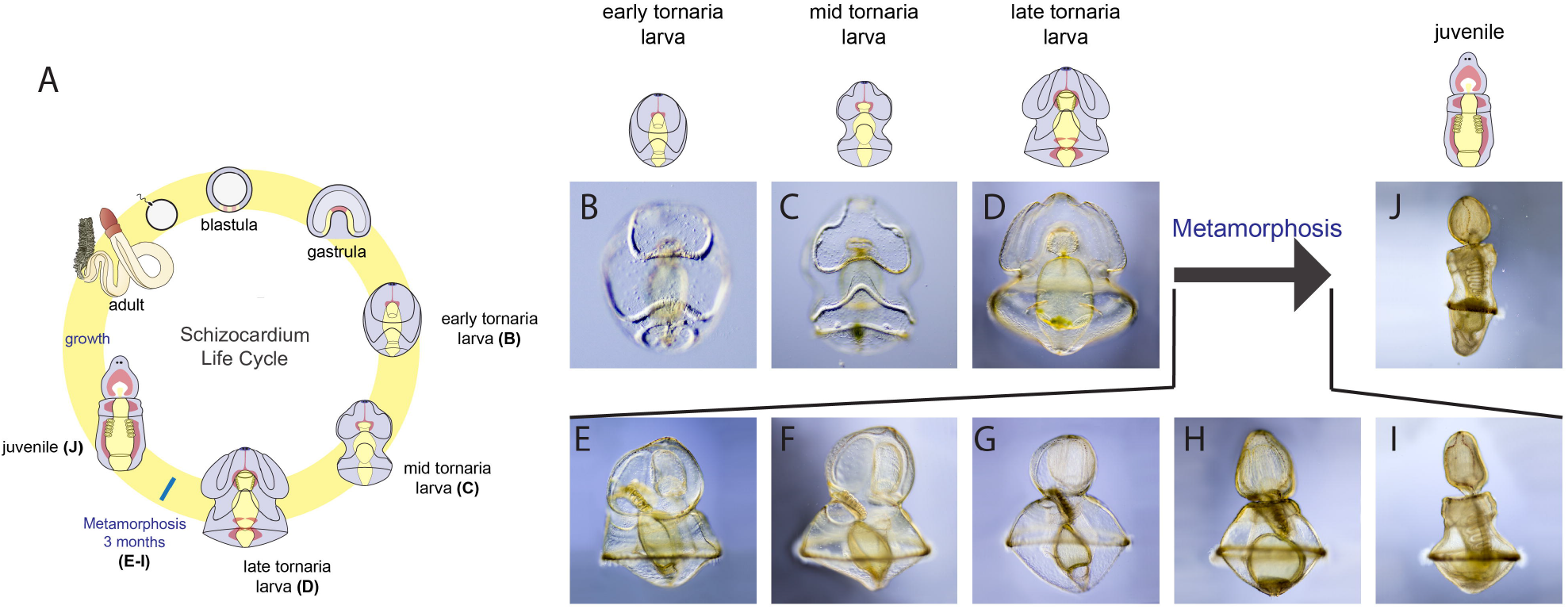
Larval development and metamorphosis of S. californicum. A) Schematic of the complex life cycle of the indirect developing *S. californicum* (modified from (34). B-J) Light microscopy of the complex life cycle of the indirect developing *S. californicum* (from (34)). B) Early tornaria larva, C) Mid tornaria larva, D) late tornaria larva, E-I) Process of metamorphosis, J) juvenile.

## Results

### Patterns of proliferation in larval and adult body plans

In an organism with a distinct larval and adult we wanted to test whether patterns of cellular proliferation involved in the development of the larva were similar or different to those during the development of the adult: is proliferation broadly distributed or detected in specific proliferative zones, and is this different between life history stages? In order to describe the distribution of proliferative cells throughout the development of *S. californicum,* we assessed the incorporation of the thymidine analog 5-ethynyl-29-deoxyuridine (EdU), which labels cells in S phase (52), at a range of developmental stages; during early larval development (Figure 1B), mid larval development (Figure 1C), late larval development (Figure 1D), during metamorphosis (Figure 1E-I), and in juvenile development (Figure 1J).

The earliest larval developmental stage consists of a tightly packed ciliary band that loops around the larva, a thin wide squamous epithelium, apical tuft, and tripartite gut (Figure 1B, Figure 2). On the ventral surface we detected EdU^+^ cells throughout the preoral and postoral loops of the circumoral ciliary band (Figure 2B, Figure 2C). The ciliary band is used for both swimming and particle capture at this stage (53, 54) and makes up a large percentage of ectoderm cells. We tested whether the ciliary bands are more proliferative than the general ectoderm, or simply have higher cell densities. The ciliary bands are densely packed with nuclei, there are ∼59% greater number of cells in the ciliary bands versus all other tissues (paired t-test, p = 0.016) and they are also more proliferative with ∼22% higher number of Edu^+^ cells vs. all other tissues (paired t-test, p=0.008) (Supplement 1A, Supplement 1B). This suggests that while the ciliary bands are nuclei-dense regions, they appear to be some of those most proliferative structures at this stage. This pattern aligns with what has been observed in the ciliary bands of other Ambularianans such as the bipinnaria larvae of *Pisaster ochraceus* and *Patiria miniata* (55). On the dorsal side of the larva in the most anterior regions, EdU^+^ cells are detected around the apical organ (Figure 2E, Figure 2F) which is the most prominent structure of the larval nervous system (56–58). Other important proliferative structures of larva include the digestive tract where microalgae that have been captured by the ciliary bands are passed from the mouth into the pharynx, and finally into the stomach, where they are digested (Figure 2H, Figure 2I).

**Figure 2.**
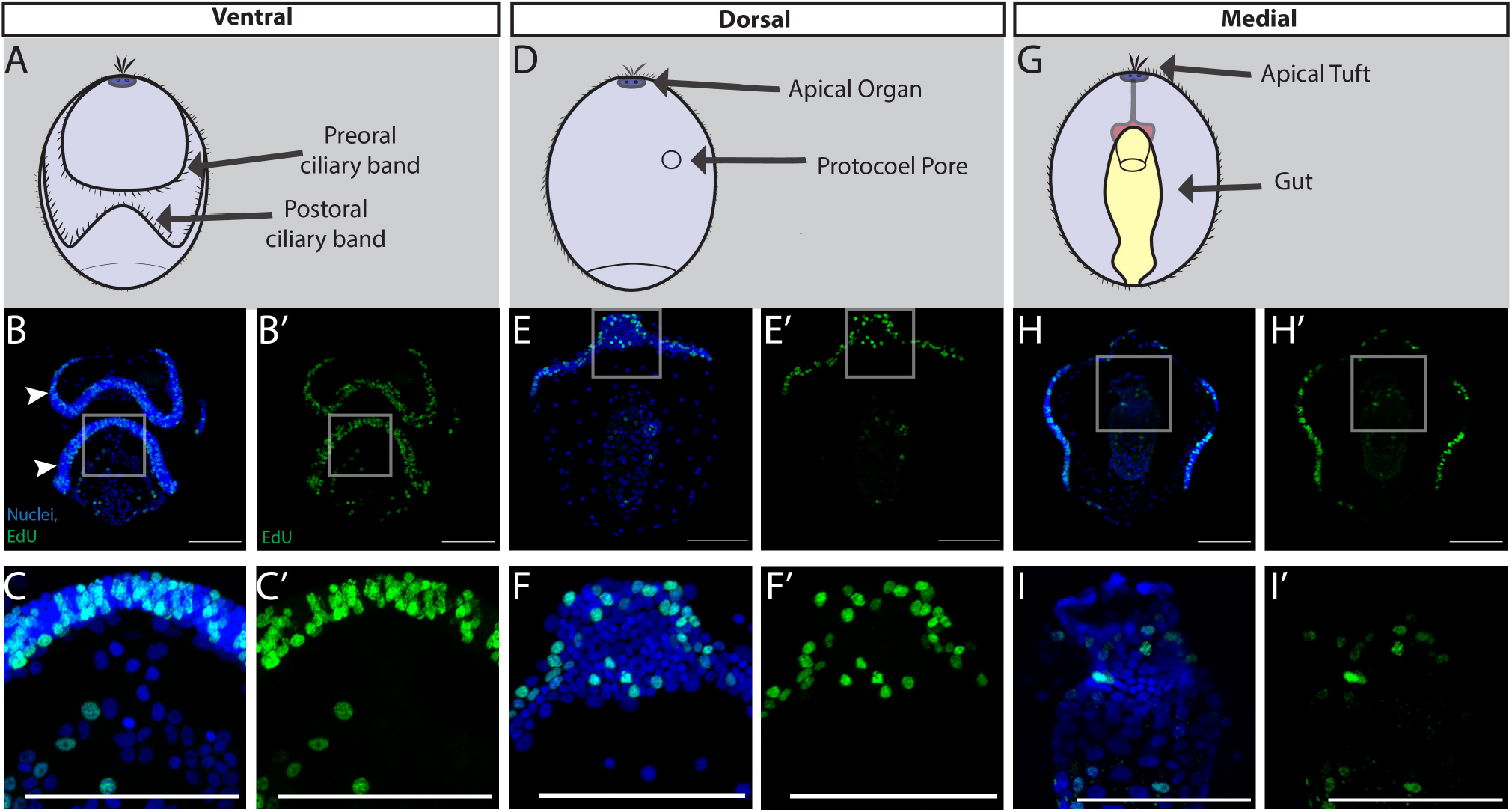
Cell proliferation throughout early larval development of S. californicum. All: anterior up; blue = Hoechst, green= EdU; scale bar is 100um. EdU staining with maximum intensity projections showing ventral (B), dorsal (E), and medial H) sections. C-I) insets. White arrowheads denote preoral ciliary band and postoral ciliary band.

As the tornaria continues to grow and reaches the middle of larval development (Figure 1C, Figure 3), which is defined by dorsal and ventral saddles, as well as the clear emergence of the posterior telotroch, proliferative cells continue to be numerous throughout the ciliary bands. This is most apparent ventrally in the preoral and post-oral ciliary bands (Figure 3B). Proliferative cells are now detected where the posterior locomotory ciliary band, the telotroch, is developing (Figure 3C). The telotroch is one of the most distinctive structures of the hemichordate tornaria with long compound cilia that beat to propel the larva through the water (54). On the dorsal surface, the protocoel pore is proliferative at this stage (Figure 3E, Figure 3F). This structure is a portion of the larval protonephridial system, an excretory system that uses a cilia-driven flow for ultrafiltration of coelomic fluid from the protocoel (59, 60). Finally, at this stage the last structure of note is the tripartite gut, composed of pharynx, stomach and intestine, which continues to proliferate and grow (Figure 3H, Figure 3I).

**Figure 3.**
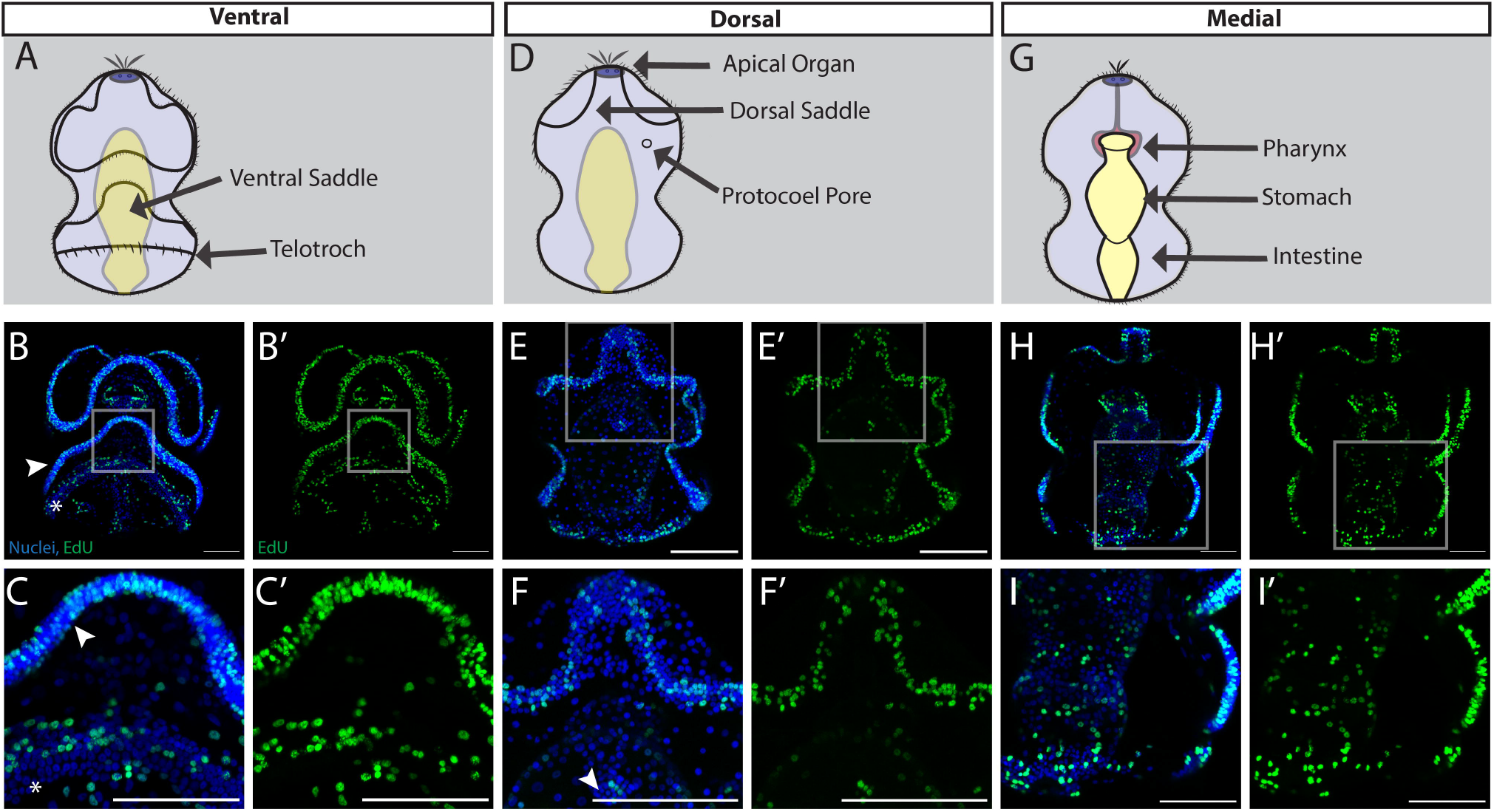
Cell proliferation throughout mid larval development of S. californicum. All: anterior up; blue = Hoechst, green= EdU; scale bar is 100um. EdU staining with maximum intensity projection showing ventral (A), dorsal (D), and medial (G) sections. C-I) insets. White arrowhead in B and C denote postoral ciliary band, white asterisk denotes telotroch. White arrowhead in F denotes protocoel pore.

Close to metamorphosis, the tornaria larva reaches full size (∼3mm) and forms two additional coelom pairs, the mesocoels and metacoels, and the precursors to the gill slits (Figure 1D, Figure 4), and we observe a notable shift in proliferative patterns from earlier developmental stages. Proliferative cells are still distributed throughout the ventral ectoderm, both in the ciliary bands, but now also more broadly in the squamous epithelium between the ciliary bands (Figure 4B). There are also a number of EdU^+^ cells distributed broadly throughout the posterior ectoderm of the larva, which is a territory that undergoes large changes at metamorphosis (Figure 4C). Across the dorsal surface of the late larva, there are numerous proliferative cells distributed throughout the epithelium (Figure 4E). There are EdU^+^ cells throughout the telotroch and on either side of the dorsal midline where the dorsal cord is beginning to form (Figure 4F). Perhaps most interestingly, at this stage, the gut has stopped proliferating and EdU^+^ cells are now detected in forming adult structures (Figure 4H). In particular, we see EdU^+^ cells in the anlage of the gill slits, which are a prominent endomesoderm feature of the juvenile body plan that are not yet functional in the late larva (61) (Figure 4I). EdU^+^ cells are also enriched in the single anterior protocoel (Figure 4J), and more posterior paired mesocoels (Figure 4K) and metacoels (Figure 4L), which will later form the adult mesodermal derivatives of the proboscis, collar, and trunk respectively. In line with previous morphological observations, in late larvae, structures of the juvenile body plan begin to proliferate to build the adult anatomical structures ahead of metamorphosis (34).

**Figure 4.**
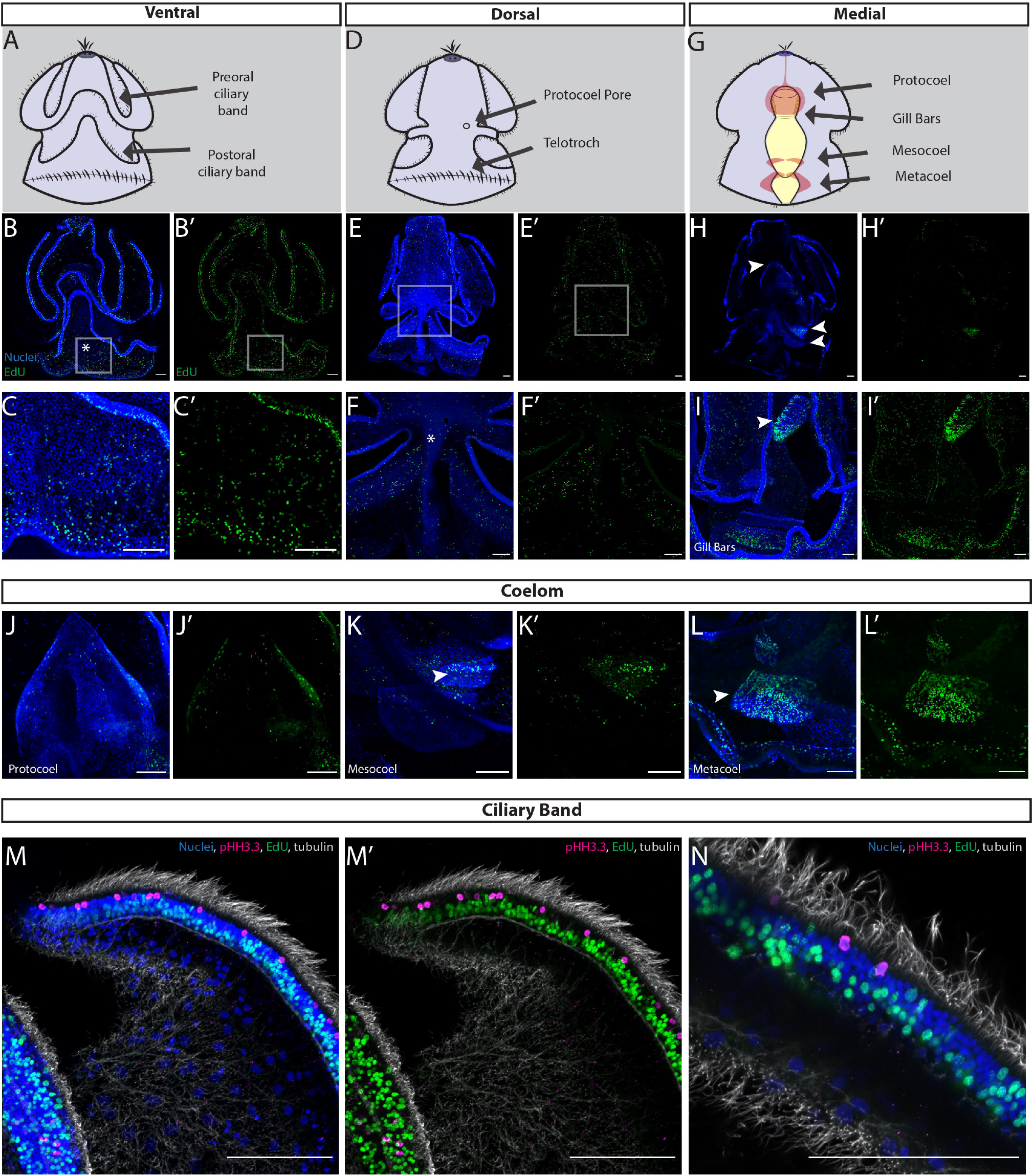
Cell proliferation in late larval development of S. californicum. All: anterior up; blue = Hoechst, green= EdU; scale bar is 100um. EdU staining with maximum intensity projection ventral (B), dorsal (E), and medial (H) sections. B) late larva ventral surface, C) inset of B, highlights ventral posterior epidermis and ciliary band E) late larva dorsal surface, F) inset of E highlights dorsal cord, marked by white asterisk, H) late larva medial section, arrowheads highlight regions that will give rise to protocoel, mesocoel, and metacoel. I) inset of lateral view of late larva medial section, arrowhead highlights gill bars. J) protocoel, mesoderm that will form the proboscis, K) mesocoel, mesoderm that will form the collar, L) metacoel, mesoderm that will form the trunk. M,N) distribution of anti-histone H3 (phospho S10) and EdU positive cells in ciliary bands, magenta= pHH3.3, grey= acetylated tubulin.

We also looked in more detail at the proliferative patterns in the ciliary band, one of the most distinct and proliferative structures of the tornaria larva (Figure 4M, Figure 4N). To achieve this, we coupled our EdU detection with immunofluorescence staining of acetylated tubulin to visualize cilia and phosphorylated serine 10 of histone H3 (pHH3.3) which marks cells in G2/M phase. We found that proliferative cells display distinct spatial distribution with a row of EdU^+^ cells at the base, then a row of differentiating phosophohistone h3.3 cells that are lateral to the cilia (Figure 4M, Figure 4N). This regional localization of EdU^+^ cells in relationship to the differentiating phosophohistone h3.3 cells suggests that there may be a specific population of proliferative cells that give rise to the ciliary bands.

### Proliferative patterns shift at metamorphosis

The first morphological indication of the onset of metamorphosis in *S. californicum* is a compaction and reorganization of the larval epidermis and an expansion of all the coeloms which results in a decrease of the blastocoelar space (Figure 1E, Figure 1F) (34). Early in metamorphosis, the primary ventral lobe and primary dorsal lobe compact around the lateral food groove (62) and EdU^+^ cells are distributed throughout several regions of the ectoderm (Figure 5B). EdU^+^ cells are distributed in the postoral field and primary dorsal saddle that give rise to both the proboscis and also around the thickening collar (Figure 5B). At this stage, EdU^+^ cells are also found in the collar and posterodorsally in the region of the developing dorsal cord (Figure 5C). On the ventral surface, EdU^+^ cells show a similar distribution to the dorsal side with proliferative cells in the pre-oral field, around the collar, in the anlage of the gill slits and in the epidermis where the ventral cord will eventually form (Supplement 1C).

**Figure 5.**
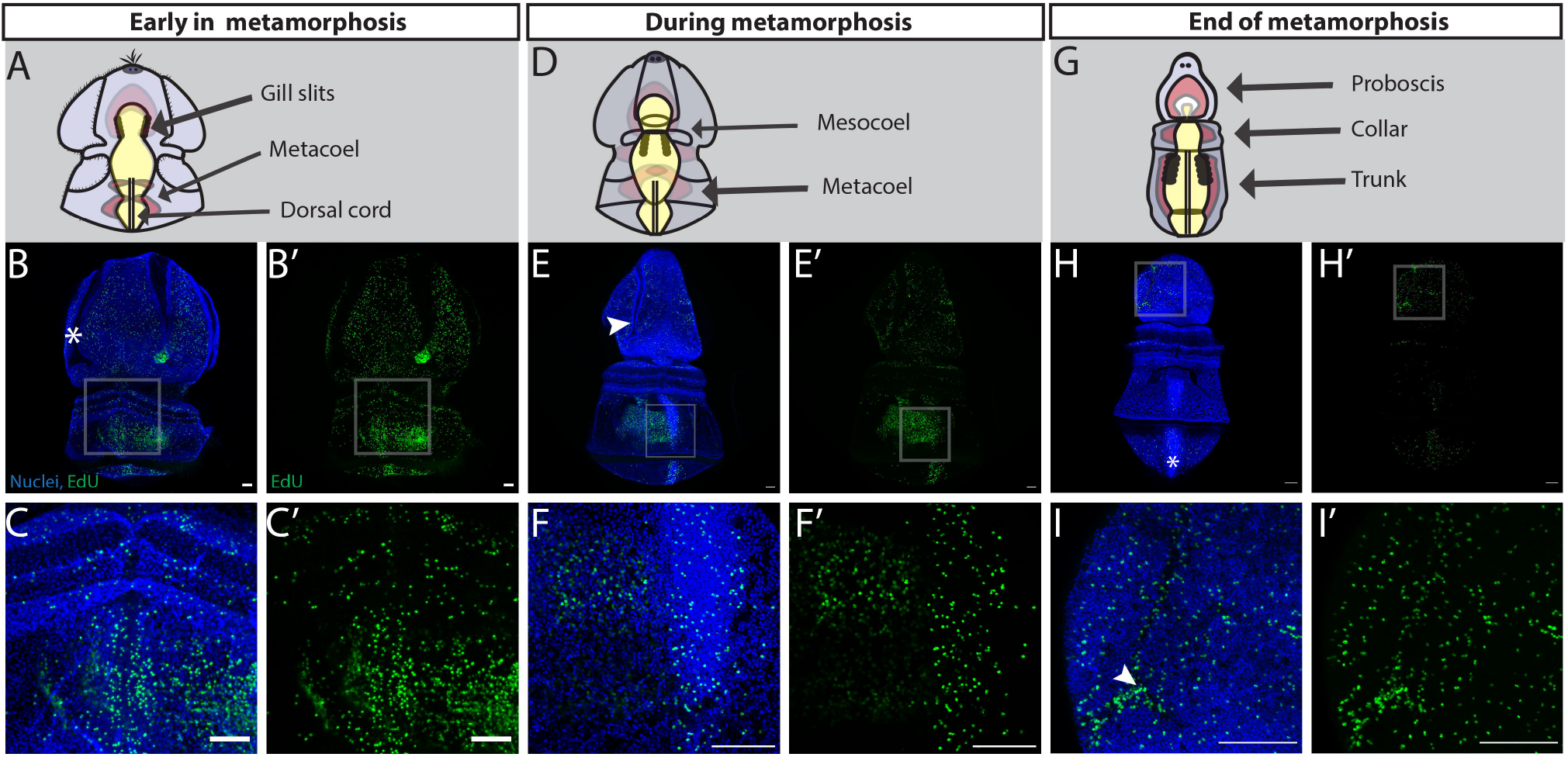
Cell proliferation throughout the metamorphosis of S. californicum. All: anterior up; dorsal view; blue = Hoechst, green= Edu; scale bar is 100um. EdU staining with maximum intensity projection: B) Early in metamorphosis, white asterisk marks lateral food groove C) Inset of B, EdU positive cells are distributed around and in the dorsal cord. E) Middle of metamorphosis, white arrowhead marks lateral food groove. F) Inset of E, EdU positive cells are found in the dorsal cord and mesocoel. H) End of metamorphosis, asterisk marks dorsal cord. I) Inset of H, white arrowhead highlights EdU positive cells are distributed throughout the lateral grooves.

Metamorphosis then proceeds with the prospective proboscis ectoderm continuing to thicken as the blastocoel shrinks bringing it in contact with the expanding anterior coelom (Figure 1G, Figure 1H). The posterior ectoderm continues to expand as the forming trunk continues to elongate. At this stage, ectodermal proliferation continues in the general epidermis of the proboscis but is absent from the remnants of the ciliary bands (Figure 5E). The epidermis of the proboscis transforms into a columnar organization as the larva begins to take on a more vermiform shape. Other proliferative regions at this stage include the developing gill slits, the metacoel, and the dorsal cord (Figure 5F). On the ventral surface at this stage proliferation occurs in the anterior ectoderm, similar to the dorsal surface, absent from where the ciliary bands had been (Supplement 1D). There are also EdU^+^ cells detected around the collar and around the field where the ventral cord will form (Supplement 1D).

Finally, metamorphosis concludes as the blastocoelar space of the proboscis is eliminated bringing the mesoderm and ectoderm in direct contact, the ectoderm of the proboscis and collar transformed into a columnar epithelium, and the posterior coeloms expand and differentiate as the trunk is elongating and narrowing (Figure 1I). At this stage, we detect proliferative cells e specifically in the proboscis, the collar, dorsal cord, and more broadly below the telotroch in the most posterior ectoderm and mesoderm (Figure 5H). Specifically, in the proboscis there are both EdU^+^ cells distributed throughout but also a clear enrichment of EdU^+^ cells in the lateral groove, the region that had previously been the larval food groove (Figure 5I). A lateral view at this stage in metamorphosis, highlights cell proliferation in the gill slits and gut as well as the dorsal and ventral midlines that give rise to the nerve cords (Supplement 1E).

In three main regions of the newly formed juvenile (Figure 1J), there is cell proliferation detected in the proboscis, collar, gill pores, gill bars, and trunk (Figure 6B). In the anterior of the juvenile, EdU^+^ cells are localized to the epidermis and line the lateral groove and anterior collar (Figure 6C). This region of the animal is highly innervated as the newly forming worm begins to burrow and explore the substratum. Proliferative cells are also found in the dorsal gill pores which have perforated to allow water to flow out and over the gill slits (Figure 6D). Finally in the posterior of the newly formed juvenile, proliferative cells are located running along the dorsal cord of the trunk (Figure 6E). At this stage on the ventral surface, we detect EdU^+^ cells in the proboscis, the gill slits, and now in the ventral cord (Supplement 1F). To see if these patterns of juvenile growth continue well after metamorphosis, we grew animals in sand for several weeks and repeated the EdU^+^ labeling and cleared the tissue to make it possible to visualize the distribution of proliferation in larger, thicker tissue. In continued juvenile growth (Figure 6F) proliferative cells can be found at the base of the collar, a region where serotonergic neurons are found (34). Interestingly, at this later stage there is no longer a strong enrichment of proliferation in the gill slits or dorsal cord but instead proliferative continue down the trunk, potentially a region of posterior growth (63).

**Figure 6.**
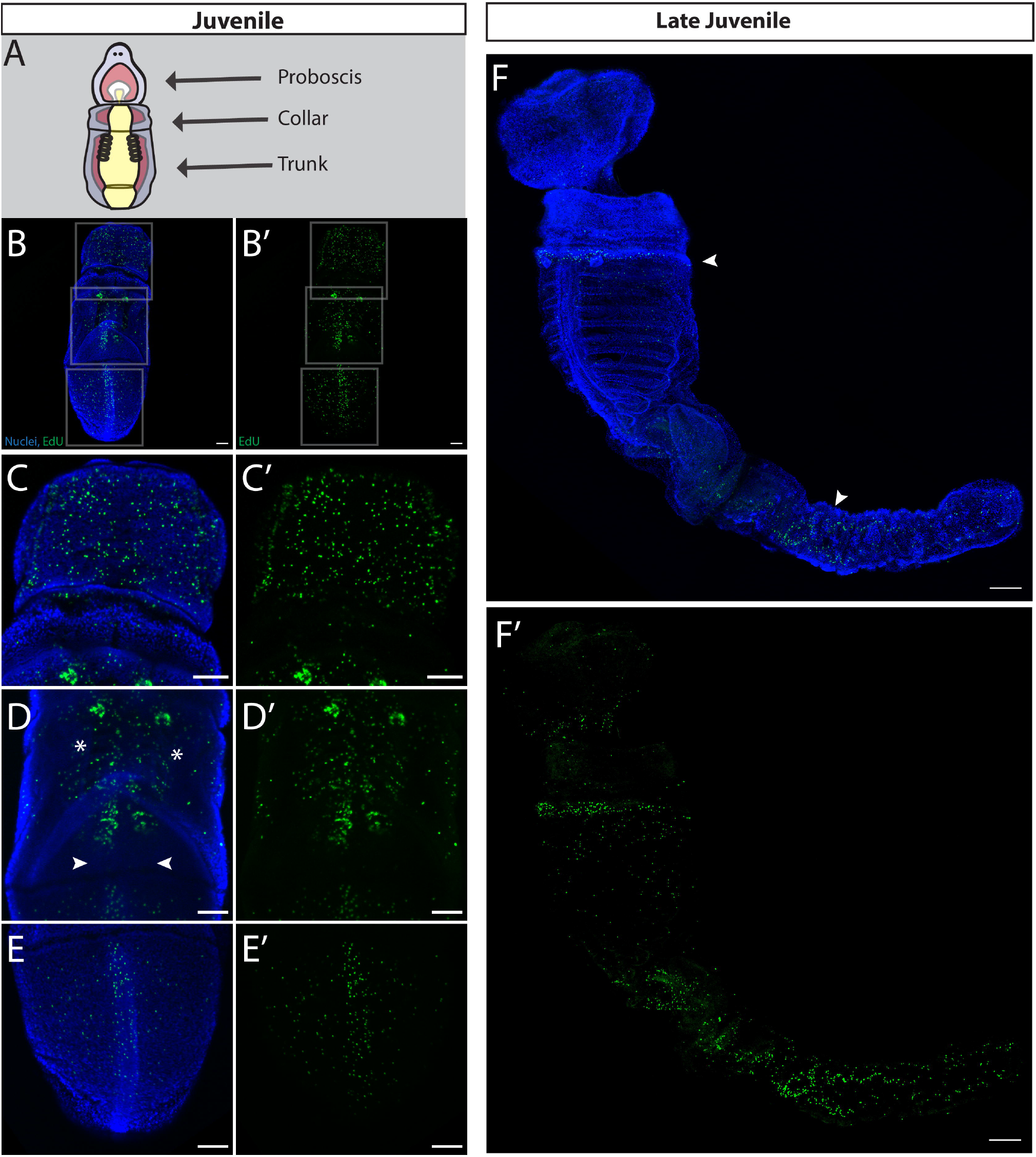
Cell proliferation in juveniles of S. californicum. All: anterior up; dorsal view; blue = Hoechst, green= Edu; scale bar is 100um. EdU staining with maximum intensity projection: B) End of metamorphosis. C) Highlights regions of B, EdU positive cells are distributed throughout the proboscis. D) Highlights regions of B, EdU positive cells are distributed throughout the gill bars. E) Highlights regions of B, EdU positive cells are distributed throughout the dorsal cord. F) Continuing development of the juvenile body plan. Arrowheads mark the base of the collar and expanding trunk

As *S. californicum* transitions from a distinct larva through metamorphosis and transforms into the juvenile, proliferative cells shift in their distribution, restricting to specific regions in the juvenile body. Overall, our data suggests that the proliferation of the adult body plan begins at late larval stages *prior* to the start of the metamorphosis itself.

### RNAseq after irradiation reveals the genetic signature of proliferative cells in two distinct life history states

To further explore the molecular characteristics of proliferative cells in *S. californicum*, we exploited the sensitivity of proliferative cells to irradiation (64–68). We hypothesized that the transcripts of irradiation-sensitive genes would be restricted to our EdU^+^, proliferative cell population. When we inspected the morphology of EdU^+^ proliferative cells with fluorescent in situ hybridization (FISH) to detect histone h2b messenger RNA, a known cell cycle gene, we find that EdU^+^ cells possess a narrow rim of cytoplasm of h2b mRNA surrounding their nucleus and these cells often display a cytoplasmic projection (Supplement 1G, Supplement 1H). This morphology is reminiscent of the proliferative cells studied in other organisms, such as planarian neoblasts, which have been characterized as rounded mesenchymal cells with a high nuclear-to-cytoplasmic ratio that often extend a cytoplasmic projection (69, 70). With these additional characterizations we next wanted to know if these proliferative cells might share any core genetic signature with proliferative cells in other organisms. One hypothesis was that *S. californicum* would have a stem-cell-like population that expresses many of the classic multipotency or germline multipotency factors, such as *piwi, vasa, nanos*, (66, 71)

To do this, we treated larvae and juveniles with irradiation. Three days after treatment animals looked morphologically the same as controls, but EdU incorporation was eliminated in both larvae and juveniles (Figure 7A, Figure 7B, Figure 7C, Figure 7D). We extracted total RNA from this same stage of three days post-irradiation from 5 pooled individuals in three independent biological replicates and made RNA sequencing libraries (Nugen-Tecan Genomics). RNAseq analysis of irradiated versus non-irradiated identified 20 genes in larvae and 123 genes in juveniles showing significant differential expression (log2 fold change ≥ −2) and p-adjusted value ≤ 10-6 juveniles (Figure 7E, Figure 7F), with 5 genes that were downregulated at both stages. 20 candidate genes were specific to the larval stage, including *fgfr-B* (fibroblast growth factor receptor B) and a number of genes involved in cell division such as *ince (*inner centromere protein) (72) *aspm-1* (abnormal spindle microtubule assembly) (73–75), and *dlgp5 (*disks large-associated protein 5) (76, 77) (Supplement 2A). In juveniles, 123 genes showed significant differential expression including a potential germline marker *spne-2 (*spindle-E), genes involved in proliferation such as *anln* (anilin), and genes related to a potential immune response *traf2* (TNF receptor-associated factor 2), *tlr2* (Toll-like receptor 2), and *tlr6* (Toll-like receptor 6) (Supplement 2B).

**Figure 7.**
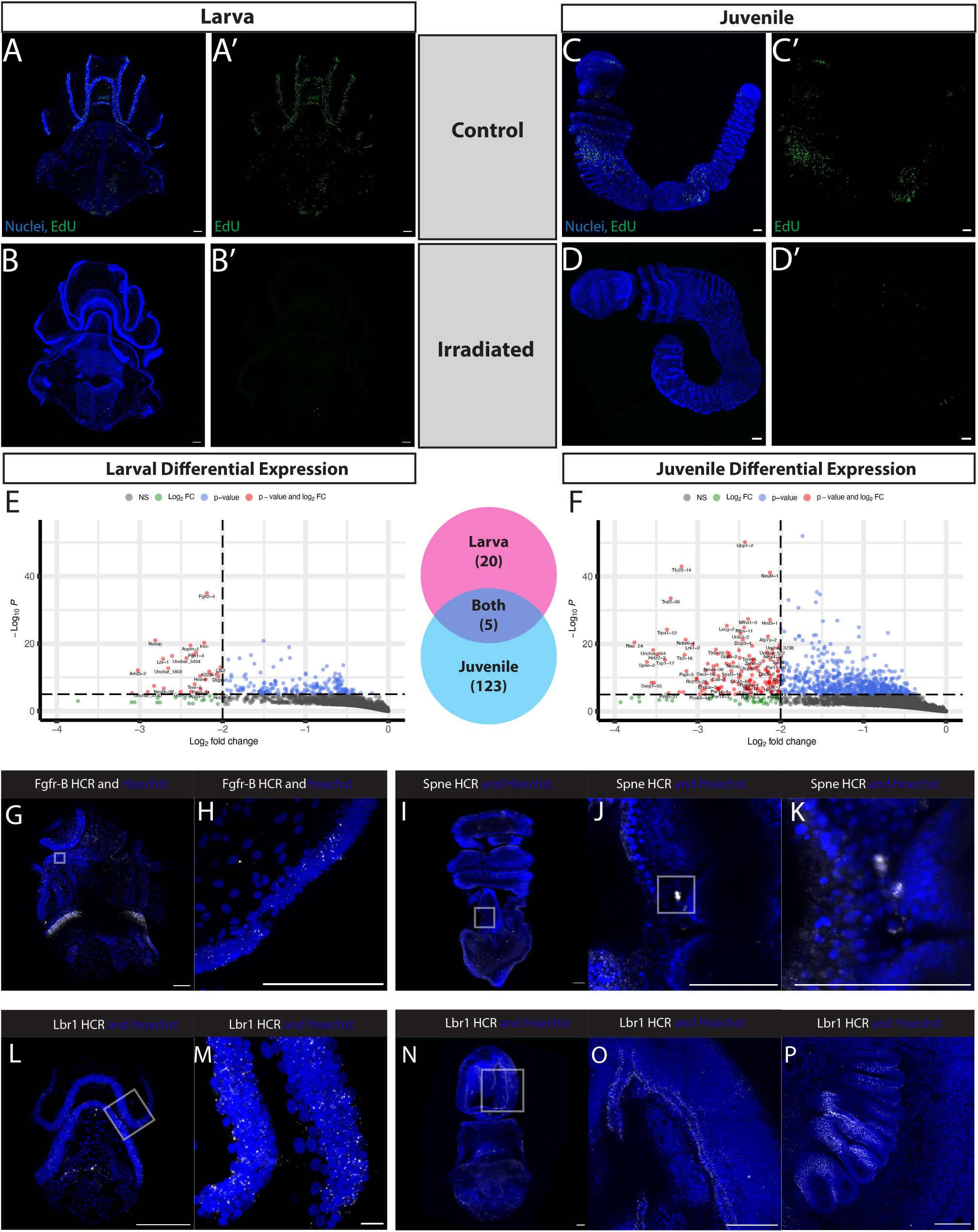
Genetic signature of irradiation sensitive EdU cells in both larvae and juveniles. All; blue = Hoechst, green= EdU; scale bar is 100um. A) Control larva representing the normal EdU pattern at this stage, representative of 5/5 animals. B) Experimental larva representing the EdU pattern at this stage after receiving 120Gy of x-ray irradiation, representative of 5/5 animals. C) Control juvenile representing the normal EdU pattern at this stage, representative of 4/4 animals. D) Experimental juvenile representing the EdU pattern at this stage after receiving 200Gy of x-ray irradiation, representative of 2/2 animals. E) Volcano plot showing expression differences in control versus irradiated larva. n=3 for each group. F) Volcano plot showing expression differences in control versus irradiated juvenile. n=3 for each group. G) A larva with HCR probes for Fgfr-B. H) Higher magnification of ciliary band with Fgfr-B transcripts. I) A juvenile with HCR probes for Spindle-E. J-K) Higher magnification of Spindle-E transcripts expressed between the ectoderm and the gills bars. L) A larva with HCR probes for Lbr-1. M) Inset of H with Lbr-1 transcripts distributed throughout the ciliary band. N) A juvenile with HCR probes for Lbr-1. O) Inset of I with Lbr-1 transcripts distributed throughout the lateral grooves. P) Lbr-1 transcripts distributed in the gill bars.

Finally, 5 genes with differential expression were shared between larva and juvenile: *lbr-1* (Lamin B Receptor), *nusap* (Nucleolar And Spindle Associated Protein 1), *tenr-5* (Tenascin-R), *tlr6-1 (*Toll-like receptor 6), and *unchar_4293* (an uncharacterized gene). *nusap* plays a role in spindle microtubule organization and also has been implicated in WNT signaling and metastasis (78)*, and tenr-5* (Tenascin-R), belongs to a group of extracellular matrix proteins, tenascins, which are important in vertebrates stem cell niches for tissue formation, cell adhesion modulation, and the regulation of proliferation and differentiation (79).

Among the larval irradiation-sensitive transcripts, *fgfr-B* was most notable. FGF receptors in vertebrates are known to regulate cell proliferation, differentiation, as well as playing a key role in pluripotent stem cells (80). The two hemichordate FGF receptors Fgfr-A and Fgfr-B arose from a hemichordate-specific duplication (81) and in the direct developing hemichordate *Saccoglossus kowalevskii, fgfr-B* is expressed in the endomesoderm at early gastrula stage and also in the ectoderm beginning at late gastrula into later stages while *fgfr-A* is also expressed (82). In *S. californicum*, we examined the distribution of *fgfr-B* mRNA and found expression throughout regions where we also had previously observed EdU^+^ cells, particularly in the ciliary bands (Figure 7G, Figure7H).

In the juvenile transcriptomes, the differential expression of *spne-2 (*spindle-E) was most notable. In *Drosophila melanogaster* spindle-E is involved in the generation of germ cell piwi-interacting-RNAs (piRNAs) and the DExD-box helicase domain of s*pindle-E* is required for silencing of transposable elements in the germline (83, 84). In *S. californicum, spindle-E* was specifically expressed in mesenchymal cells around the posterior of the gills bars (Figure 7I, Figure 7K), which is consistent with Vasa expression in *P. flava* (85). Given that *spindle-E* was expressed in a similar region and has been implicated in germline regulation, we hypothesize that *spindle-E* could potentially be a marker of proliferative germline cells in hemichordates.

Of genes that were differentially expressed in both life histories, of particular interest is the lamin B receptor gene, *lbr-1*, which plays an important role in tethering chromatin and maybe be a universal mark of proliferative cells across both life history stages of *S. californicum*. There are two types of chromatin attachment to lamina, one type is executed by the lamin b receptor in embryonic and non-differentiated cells, and the other by specific lamin a/c binding proteins in differentiated cells (86). *lbr-1* orthologs have previously been identified in ascidians and echinoderms and previous work has suggested that this gene may have emerged with the appearance of the deuterostomes (87). We examined the expression of *lbr-1* in larvae and found it localized in the ciliary bands (Figure 7L, Figure 7M), which we previously demonstrated were regions of active cellular proliferation (Figure 3A). Similarly, in the juvenile stage we found *lbr-1* expression in a similar territory where we had observed the distribution of EdU^+^ cells (Figure 5F, Figure 6B, Figure 6C), such as the lateral grooves in the proboscis (Figure 7N, Figure 7O) and in the gill bars (Figure 7P). Our findings suggest that expression of *lbr-1* might serve as a useful marker of labeling proliferative cells across both life history states.

Finally, the classic multipotency or germline multipotency factors, such as *piwi, vasa, nanos*, (66, 71) not have significant differential expression (Supplement 2C, Supplement 2D). Instead, what we recovered from our study were genes related more to specific proliferative populations (*fgfrB, spne-2,* and *lbr-1)* and thereby, revealed a possible heterogeneity among proliferative progenitor cells.

### Cell death remodels larval tissue at metamorphosis

After an investigation of cell proliferation throughout the life cycle and metamorphosis of *S. californicum,* we next turned our attention to cell death to determine whether the distinct patterns of proliferation between life histories are also correlated in patterns of cell death. One larval structure that is lost or extensively remodeled at metamorphosis, is the circumoral ciliary band, also called the longitudinal ciliary band, a larval specific feeding structure (57) that is not retained in the juvenile. We investigated the distribution of cell death with TUNEL (terminal deoxynucleotidyl transferase dUTP nick end labeling), which detects breaks in DNA as a proxy for cells undergoing apoptosis (88). We overcame previously limitations of TUNEL detection by taking advantage of Click-iT technology, which utilizes a modified dUTP with a small, bio-orthogonal alkyne moiety (EdUTP) and a copper catalyzed covalent reaction click reaction between that alkyne and a picolyl azide dye (89, 90).

Throughout larval development and in late larva we detected few TUNEL^+^ cells, suggesting very limited cell death at larval stages (Supplement 3A, Supplement 3B, Supplement 3C). However once metamorphosis begins, as indicated by the thickening of the larval epithelium, there is a large increase in TUNEL^+^ cells (Figure 8B). TUNEL^+^ cells are distributed broadly throughout the ectoderm, with most of them on either side of the developing dorsal cord, and in and around the circumoral ciliary band (Figure 8C). The circumoral ciliary band is labelled with a large number of TUNEL^+^ cells, supporting the morphological observation that this structure begins to break down at this stage. There also are a small number of TUNEL^+^ at the anterior end of the protocoel (Supplement 3D, Supplement 3E). TUNEL^+^ cells are also absent from any part of the gut at this stage, which is consistent with the observation that the gut is maintained throughout the transition from larvae to adult (34). In order to confirm adequate penetration of the TUNEL labeling into the deeper tissue layers, we performed a positive control by artificially nicking the ends of DNA with DNAse-1 (Supplement 3G, 3H).

**Figure 8.**
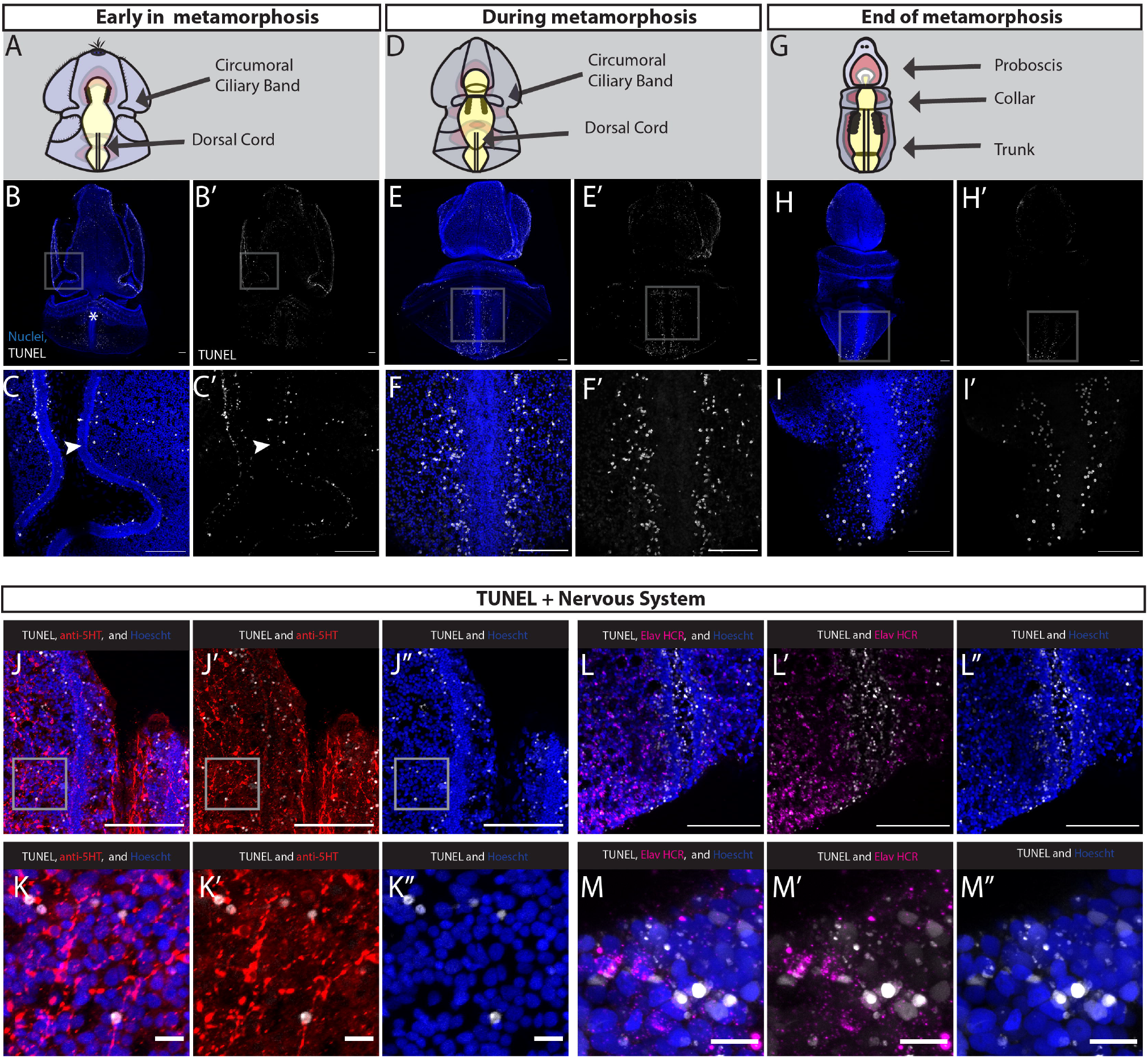
Cell death throughout the metamorphosis of S. californicum. All: anterior up; blue = Hoechst, grey= TUNEL; scale bar is 100um. B) Start of metamorphosis with an increase in TUNEL^+^ cells. C) Highlights regions of B), specifically around lateral grooves. E) Middle of metamorphosis. F) Highlights regions of E), specifically around the dorsal cord. H) End of metamorphosis. I) Highlights regions of H), specifically around the dorsal chord and collar. J) Overlap of serotonin^+^ cells and TUNEL^+^ cells. K) Inset of G, highlighting TUNEL^+^ and serotonin^+^ positive cells. L) Overlap of Elav^+^ cells and TUNEL^+^ cells. M) Inset of L, highlighting TUNEL^+^ and Elav^+^ cells.

At the mid-metamorphosis stage, TUNEL^+^ cells are distributed throughout the epidermis and continue to label the disintegrating ciliary bands that begin to fuse with each other (Figure 8E). At this stage we continue to detect very few TUNEL^+^ cells in the mesoderm and endoderm. The other region of the ectoderm where the greatest number of TUNEL^+^ cells are found is directly lateral to the dorsal nerve cord (Figure 8F), a region where we observed a large number of proliferative cells at the same stage. These general patterns that we see on the dorsal side are broadly similar to what is observed on the ventral surface with TUNEL^+^ cells distributed in the ciliary bands and broadly in the ectoderm but excluded from the ventral nerve cord (Supplement 3F). The presence of TUNEL^+^ cells in and around the ciliary bands is consistent with a previous observation that during metamorphosis the circumoral ciliary band degenerates, and the serotonergic nervous system in this region undergoes extensive reorganization, as neurite bundles that are present in the ciliary grooves disappear as the ciliary bands fuse (34, 91).

Finally at the end of metamorphosis, there are fewer TUNEL^+^ cells detected. In the anterior, there are very few TUNEL^+^ cells, the proboscis ectoderm is compacting as the mesodermal protocoel has finished expanding (Figure 8H). Those TUNEL^+^ cells that remain are scattered throughout the epidermis in the proboscis, but no longer in the lateral grooves, which from our EdU study we have observed becoming proliferative at this stage (Figure 5F). In the posterior, the remaining TUNEL^+^ cells are detected on either side of the dorsal cord, most prominently in the posterior where the larval epidermis is still collapsing (Figure 8I).

To understand the interaction of cell death with the nervous system, we examined serotonin localization along with TUNEL early in metamorphosis and found there is colocalization of serotonergic^+^ cells with TUNEL^+^ cells (Figure 8J, Figure 8K). We also examined expression of Elav, a pan-neuronal marker, with TUNEL (Figure 8L, Figure 8M) and found a number of colocalized Elav^+^ and TUNEL^+^ cells at the edge of the epidermis. Our findings suggest that portions of the larval nervous system undergo cell death at metamorphosis and that the nervous system of the anterior ciliary bands may not be maintained in the juvenile body plan.

Overall, from our characterization of cell death, we find that an increase in TUNEL^+^ cells correlated with metamorphosis. We found TUNEL^+^ cells broadly distributed in the epidermis and that their restriction over time, from anterior to posterior, correlates with the morphological observation of an anterior to posterior temporal progression of ectodermal thickening (34). While we cannot rule out additional forms of tissue remodeling or histolysis, our findings suggest that cell death plays an important role in remodeling larval structures specifically in the anterior ciliary bands, which fuse during metamorphosis.

## Discussion

The development of an adult by transformation of a larva is very common in bilaterians, yet we understand very little about the details of how this process occurs at a cellular level, particularly given how the process of metamorphosis differs across groups with different life history strategies (10,92–94). This study in the hemichordate *S. californicum* focuses on characterizing the patterns of cellular proliferation and cell death during the two different life history stages and during metamorphosis. Unlike other certain model species like *D. melanogaster*, where metamorphosis results in a major histolysis of larval tissues and the adult emerging from imaginal discs (95), morphological studies in *S. californicum* suggest that metamorphosis occurs by remodeling of larval tissues and the transformation of larva into the adult (34). We have investigated similarities and differences between the life history stages in patterns of proliferation and cell death to investigate their contribution to how and when larva and adult body plans form. Our work illustrates that the development of the distinct life history states in *S. californicum* occurs via a combination of regionalized cell proliferation and limited cell death, and that suggest that metamorphosis represents a transformation of a larval body plan rather than building the adult de novo from a small population of cells.

### The larval body plan shaped by proliferation

For an organism with indirect development, rapid growth of the larva is essential. Eggs are small, yet juvenile size at metamorphosis is a good indicator of individual fitness, so larval growth before metamorphosis is critical (12, 96). We observe in early larval development that the tornaria larva is formed primarily through cell proliferation and limited amounts of cell death (Figure 2, Figure 3, Supplement 3A, Supplement 3B, Supplement 3C). The patterns we observe highlight regional differences with proliferative cells contributing to growth: proliferative cells are distributed throughout the larval epidermis, the gut, and prominently in the ciliary bands (Figure 2A, Figure 3A). But we observed little cell death at this stage in larval development with the use of the TUNEL assay (Supplement 3A). There are some TUNEL^+^ cells distributed throughout the larva, but it does not appear that these cells are concentrated to any particular structure or tissue. While cell death in bilaterian larval development has not been surveyed broadly across taxa, cell proliferation in larval development has been assessed in a number of marine larvae and the patterns we observe in *S. californicum* confirms and extends what has been found in other species (55).

### Origin of the adult body plan begins in the late larva

The morphological discontinuity between larvae and adults has been a consistent source of great curiosity for zoologists (10,97,98). While metamorphosis is often thought of as the time when the adult animal emerges from the vestiges of larval anlage, there can often be important events that happen prior to the start of any clear morphological metamorphosis (1, 6). In the late larva of *S. californicum*, specific regions of the developing adult body start to proliferate prior to what has been described as the metamorphosis in indirect-developing hemichordate (Figure 4). This is most obvious in the proliferation of the coeloms; the protocoel, mesocoel, and metacoel that give rise to the proboscis mesoderm, collar mesoderm, and trunk mesoderm respectively (Figure 4H, Figure 4J, Figure 4K, Figure 4L). The formation of some of these morphological landmarks has been described previously (34), but our study clarifies that these structures originate via proliferation. There are other structures of the hemichordate adult that begin to form within the larva, such as the gill slits, and elements of the adult nervous system, which was also observed in another indirect-developing species (99). Adding to these morphological observations, our study has shown that these tissues and structures proliferate prior to the start of metamorphosis, and supports the hypothesis that this is essential in preparing the organism for a major life history transition (1). However, at this point in larval development little cell death is detected by the TUNEL assay as well as morphological observations (Supplement 3B, Supplement 3C). This will begin to change at metamorphosis.

### Metamorphosis integrates cell proliferation and cell death

We found that cell death correlated with the onset of metamorphosis, and regionalized cell proliferation that began during late larval development continues into the adult. Cell death is detected in regions where larval specific structures are being remodeled (Figure 8B, Figure 8C) and likely important in shaping the morphogenesis of emerging adult structures, such as the dorsal cord of the forming adult nervous system (Figure 8E, Figure 8F, Figure 8H, Figure 8I). Clearly the onset of adult morphogenesis, and the initiation of overt metamorphosis, results in a major shift in the patterns of proliferation and cell death.

Cell death within larval-specific structures has long been implicated in studies of metamorphosis, indeed one of the first recorded observation of apoptosis was in the metamorphosis of the toad, *Alytes obstetricans*, in which it was noted that cells of the notochord disappeared and were replaced by cells of the vertebrae (100). Since then, there have been important findings in the role of cell death during anuran metamorphosis that have extended the importance of timing in this process (101–103). The mechanism of metamorphosis in insects such as butterflies and fruit flies have also provided important comparative perspectives into the role of programmed cell death as a key event in this process (104, 105). Finally, cell death has been implicated in the metamorphosis of marine invertebrates as they transition from planktonic larvae to benthic juveniles (106), in particular in the remodeling of the larval nervous system in gastropods (107, 108). In *S. californicum* we begin to detect TUNEL^+^ cells at the start of the morphological metamorphosis, most obviously around the anterior ciliary bands, which are involved in larval feeding, but are broken down during metamorphosis (Figure 8B). This is similar to what has been observed in sea urchins where apoptotic cells were detected in the arms and ciliary bands of competent larvae (109–111). In *S. californicum* serotonergic neurons are associated with ciliary bands and we find TUNEL and serotonin double-positive cells suggesting one way these systems change at metamorphosis (Figure 8J, Figure 8K).

While apoptosis plays an important role in removing larval-specific structures at metamorphosis, it may also be involved in sculpting larval tissue during metamorphosis, reminiscent of digit development in vertebrates (112), the formation of leg joints and head segments in *D. melanogaster* (113). One defining feature of metamorphosis in *S. californicum* is an overall decrease in size that occurs from anterior to posterior. Our finding that cell death proceeds from the anterior to posterior in the dorsal epidermis (Figure 8E, Figure 8F) suggest this might be an important process in integrating and removing larval tissue, similar to the apoptosis observed in the mouse paw or fly larva (114). However, this would need to be tested more rigorously with more sophisticated functional approaches. Overall, while we find that cell death occurs both in larval-specific structures but also broadly in larval tissue during metamorphosis, there are also regions with more limited cell death such as the anterior ectoderm, collar ectoderm, and tripartite gut, leading to the possibility that not all larval cells die and are instead incorporated into the adult body plan.

Our characterization of cell death suggests that *S. californicum* does not have a classical “catastrophic metamorphosis”, which has commonly been attributed to some marine invertebrates with complex life cycles, where an uncoupling of larval and adult body plans occurs as the larva is discarded and the growth of the adult occurring through “set-aside cells.” This has been described in nemerteans and echinoderms (115, 116). Work by Peterson, Cameron, and Davidson focused on the sea urchin *Strongylocentrotus purpuratus* in which the juvenile grows from a small rudiment within the larval body from a population of sequestered cells they termed “set-aside cells’’ (18). This concept of “set-aside cells’’ has been highly influential, but recently has been revisited (117) and reframed in a new paradigm which presents the formation of larval and adult body plans in a broader developmental ontology of “deferred development.” Instead of a definition of cellular identity which is compared across wide evolutionary distances, the concept of “deferred development” focuses on timing with the specification and terminal differentiation of some cell populations relative to others (117). For example, the sea urchin *S. purpuratus* would fall under a category of deferred development that shifts the development of the adult into nonfunctional rudiments, while an organism like the annelid *Capitella tellata* would be in a different category where the deferral of the adult is at the level of entire tissue or organs, in this case a posterior growth zone (14, 118). This definition of “deferred development” makes comparisons between groups of organisms far more feasible — instead of trying to directly compare a specific population of cells, the comparisons are between developmental process and timing. This paradigm of “deferred development” is helpful for understanding the development of S. *californicum*, which is more similar to the delayed life history shift of marine annelids than that of sea urchins. In *S. californicum*, we observe this deferral of the adult in the late larva where proliferation of structures of the adult body plan occurs *prior* to the start of metamorphosis (Figure 4).

### Proliferative patterns of growth differ in the adult body plan, as do the markers of these proliferative populations

Juvenile *S. californicum* have distinct patterns of growth — proliferation continues to be enriched in structures that were not functionally part of the larva. The dorsal and ventral cords, gill bars, proboscis ectoderm and lateral groove are all clear examples of regional proliferation of adult structures (Figure 6B, Supplement 1F). We also looked later in juvenile development and found that proliferation was most striking in the trunk region of the animal that continues to grow (Figure 6F). This pattern of post-metamorphic growth is reminiscent of the posterior axis elongation by an extended period of posterior growth described in the direct-developing hemichordate *S. kowalevskii* (63).

Given that patterns of cell proliferation differed between larvae and juveniles, we wanted to test whether the genetic signature of proliferative cells was similar or different between the life history states, and if there were specific populations of pluripotent stem cells or broad populations of proliferative progenitors. In organisms such as colonial ascidians (119), acoels (120), flatworms (121, 122), cnidarians (123, 124) and sponges (125), adult stem cells retain the potential to produce both the germline and several somatic cell types, and it has been suggested that there may be an ancestral animal stem cell (126). We did not recover a clear pluripotent stem cell population, which contrasts organisms with clear neoblast populations like platyhelminthes and acoels (66). Genes associated with multipotency or germline multipotency also did not have significant differential expression (Supplement 4C, Supplement 4D). Instead, what we found was differential expression of genes like *lbr-1* in both larvae and juveniles, pointing towards the importance of chromatin state independent of the type of proliferative cell. It was previously suggested that the patterns of *lbr* and *lamA/C* expression in a number of different mammalian cell types may be indicative of peripheral heterochromatin tethers regulating differentiation and perhaps this is a larger uniting trend across deuterostomes (86).

When we looked at the differential expression in irradiated vs non-irradiated larvae, we found a relatively limited number of significantly differentially expressed genes such as those involved in cell division (such as *ince, asmp-1, dlgp5*) and most interestingly *fgfr-B* (fibroblast growth factor receptor B). A similar approach in the parasite *Schistosoma mansoni* found fgf receptors were downregulated in response to irradiation, and further showed that the inhibition of Fgf signaling with RNAi resulted in reduced EdU incorporation and down regulation of cell-cycle-associated transcripts (127). In juvenile animals, while we did recover the expression of what may be a germline specific marker in *spindle-E*, there were also a number of transcripts from our irradiation experiment that suggest a potential immune response, such as *traf2*, *tlr2* and *tlr6*. We conclude that the differential expression of these genes is likely related less to the depletion of irradiation-sensitive transcripts and more likely an immune reaction in response to the irradiation.

While additional work will need to be done to determine if markers such as *fgr-B* and *spindle-E* are lineage and life history-specific proliferation markers, our results certainly support the hypothesis that formation of larval and juvenile structures draws on distinct sets of proliferative populations. The strong focus on species with direct development may miss some interesting regulatory features of distinct proliferative cell populations related to the development of complex life cycles. For a fuller understanding of developmental diversity and how it has shaped animal body plans, we need both a broader phylogenetic sampling but also greater representation of complex developmental strategies.

### Integration of larval and adult body plans

Our study of the balance between cell proliferation and cell death has provided important insights into the timing of metamorphosis — that adult morphological elements proliferate prior to the start of metamorphosis, and that the onset of metamorphosis correlates with the onset of cell death. However, this study has also raised more questions, particularly the mysterious fate of the majority of larval cells that are seemingly maintained through the threshold of metamorphosis. The most provocative and exciting possibility is the potential of larval cells taking on new identities in the adult body plan. In sea urchins, which were the key example of catastrophic metamorphosis with set-aside cells, the organization of the larval epithelium is preserved as regionalized apoptosis occurs in the larval arms that are resorbed (111). Morphological studies of other echinoderms like sea cucumbers and cidaroids suggest that much of their larval epidermis is maintained into the juvenile stage and is not lost at metamorphosis (128, 129). Even in examples such as nemerteans, which are described as “maximally-indirect developers’’ (115), we now know that the cells that create the imaginal discs also contribute to the larval body (130). Even in *D. melanogaster* with its specialized imaginal disc cells, differentiated larval tracheal cells become proliferative and form the adult trachea and also adult-specific air sacs (131, 132). In the tobacco hornworm, *Manduca sexta,* differentiated cells of the larval legs contribute to the adult legs (133). Perhaps this linkage between larval and adult cells is far more common than had been previously appreciated. While genetic tools would need to be developed to study this in detail, studies of transformational metamorphosis have the potential test whether transdifferentiation of cell types occurs as part of normal ontological development in organisms with complex life histories. If this is indeed true, we are left with a tantalizing question, when many larval cells remain, how do they take on the appropriate function in the adult?

Almost all our understanding of adult development comes from direct developers where the adult body plan emerges from the embryo. Despite the prevalence and phylogenetic breadth of species that represent indirect development, we understand very little about how adult development occurs by transformation. Clearly a greater focus is needed on the range of development strategies that characterize metamorphosis in metazoan animals. Only through this broader sampling of life history strategies can we hope for a more comprehensive understanding of the developmental mechanisms responsible for adult body plan formation.

### Conclusions

Our study describes cell proliferation and cell death through the development of the indirect developing *S. californicum*. This species has distinct larval and adult body plans and a metamorphosis that is more transformational than catastrophic. Our data represent an important investigation into the common, yet understudied, bilaterian developmental strategy of formation of adult body plan by transformation of a larval body plan. Despite the prevalence of this life history strategy, we have little developmental insights into how this process occurs. Our study has also helped to reframe the question of timing of the formation of the adult and the onset of metamorphosis, uncovering that the proliferation of mesodermal adult body plan components starts prior to any clear signs of metamorphosis. Although cell death is a prominent feature of metamorphosis and adult body plan development, it is unlikely that the entire cell complement of the adult can be explained by larval cell death and proliferation of a distinct adult stem cell population. Altogether, our study establishes a cellular characterization of the formation of larval and adult body plans and transitions through metamorphosis in *S. californicum,* an important species for understanding this transformational process through the lens of cell, developmental, and evolutionary biology.

## Methods

### Collecting, spawning, and larval rearing

Adult *Schizocardium californicum* were collected in Morro Bay State Park, California, in a mudflat located at 35°20′56.7”N 120°50′35.6”W with appropriate state permitting. Animals were spawned as described in (34) with individual females transferred in bowls of filtered seawater and placed in an illuminated incubator at 24–26°C. Once hatched, larvae were transferred to 1 gallon glass jars with continuous stirring and fed larvae with a 1:1 mix of *Dunaliella tertiolecta* and *Rhodomonas lens*. Every two to four days, containers are washed, water was replaced with clean filtered seawater, and fresh algae was added. In order to grow larger numbers of animals, some larvae were placed on a continuous flow-through system by being transferred into diffusion tubes (134). Once animals began metamorphosis, they were transferred into glass bowls with terrarium sand.

### EdU labeling

Labeling and detection of proliferating cells were performed using the Click-it Plus EdU 488 Imaging Kit (Life Technologies), with the following modifications. Larva and juvenile worms were cultured in FSW supplemented with 10 μM EdU diluted from a 10 mM stock in DMSO. Unless otherwise noted, animals were pulsed for 30 minutes with EdU then fixed with 3.7% paraformaldehyde in MOPS fix buffer (0.1M MOPS, 0.5M NaCl, 2mM EGTA, 1mM MgCl2, 1X PBS) for 1h at room temperature (RT). For detection of EdU incorporation, labeled embryos were transferred to a solution of PBS and the detection was performed following the manufacturer’s protocol with an increased permeabilization time in 0.5% Triton® X-100 in PBS of 40 minutes and an increased detection time of 45 minutes.

### TUNEL Detection

Detection of apoptotic cells were performed using the Click-iT Plus TUNEL Assay for in situ apoptosis detection with Alexa FluorTM dyes, with the following modifications. Animals were fixed with the standard protocol (3.7% paraformaldehyde in MOPS fix buffer), washed twice in 1x PBS and permeabilized with proteinase-K for 15 minutes at room temperature. TdT reaction mixture was incubated for 60 minutes at 37°C and Click-iT Plus reaction was carried out for 30 minutes at 37°C.

### Antibody Labeling

Fixation and antibody labeling was performed as described previously (34). To visualize proliferative cells, we used a rabbit polyclonal anti-histone H3 (phospho S10) (Abcam ab5176) diluted 1:200 in blocking solution. To visualize cilia, we used a mouse monoclonal anti-acetylated tubulin antibody (Sigma T7451) diluted 1:400 in blocking solution. To visualize the serotonergic nervous system, we used a rabbit anti-serotonin antibody (Sigma S5545) diluted 1:300 in blocking solution. Secondary antibodies (ThermoFisher, Alexa Fluor) were added at 1:1000 dilution to the blocking solution.

### Imaging

Nuclei were stained with Hoechst 33342 (1:1000) in PBS and mounted in PBS using coverslips elevated with clay feet. For juvenile worms, samples were transferred into 50% glycerol for 30-60 minutes, then 70% glycerol for imaging. Samples were imaged on a Zeiss LSM 700 with 10X, 20X and 40X objectives. For samples larger than the field of view, maximal intensity projections from several stacks were stitched together (Fiji).

### Irradiation and transcriptional profiling

Larva and juvenile worms were exposed to 120 and 200 Gy of X-ray irradiation on a CellRad Faxitron source. Animals were cultured in FSW after irradiation for 3 days and purified total RNA was prepared from pools of 5 animals using Qiagen RNeasy. Three independent biological replicates were performed for both control and irradiated experimental groups. Individually tagged libraries for RNA-seq were prepared (Nugen-Tecan Genomics Universal mRNA-seq Kit), pooled in a single lane, and 75-bp pair-reads were generated using an Illumina HiSeq2000 at the Chan-Zuckerberg Biohub. The resulting reads were mapped to the annotated *S. californicum* genome (v2.0) using CLC Genomics Workbench (CLC Bio) and differential expression was conducted with DeSeq2. *apeglm* was used for log fold change shrinkage (135) and *v*st (variance stabilizing transformation) was used for visualization (136).

### In-situ hybridization

Samples were relaxed using 3.5% MgCl2 prior to fixation and fixed in 3.7% formaldehyde in MOPS fix buffer for 1h at room temperature (RT), washed in fix buffer, dehydrated in 100% ethanol and stored at −20°C. Genes were amplified from stage specific cDNA with random hexamers and cloned into pGEM-T Easy (Promega). Digoxygenin labeled antisense probes were synthesized using SP6 or T7 RNA polymerase (Promega). In situ hybridization was performed a combination of what has been described previously (50, 137). RNA probes were diluted to 0.1 – 1 ng/ml and hybridized overnight at 60°C and visualized using an Anti-DIG AP antibody and TSA-Cy3.

### In situ HCR version 3.0

Complementary DNA sequences specific to genes of interest were submitted to the in situ probe generator from the Ozpolat Lab (138). Gene orthology was determined by collecting sequences of interest from related species and then building gene trees. Sequences were aligned with MUSCLE (139) and trees were calculated with Bayesian inference trees using MrBayes version 3.1.2 (140) in 1,000,000 generations with sampling of trees every 100 generations and a burn-in period of 25% (Supplement 4). The sequences generated by the software were used to order DNA oligo pools (50*μ*mol DNA oPools Oligo Pool) from Integrated DNA Technologies, resuspended to 1*μ*mol/*μ*l in 50mM Tris buffer, pH 7.5. HCR amplifiers with fluorophores B1-Alexa Fluor-546, B2-Alexa Fluor-488, and B3-Alexa Fluor-647 were ordered from Molecular Instruments, Inc. The HCR was performed based on Choi et al., 2018 and the Hybridization Chain Reaction (HCR) In Situ Protocol from the Patel Lab (141, 142). For experiments that involved HCR and TUNEL labeling, HCR was conducted first, and then additional labeling was performed after.

## Declarations

### Ethics approval and consent to participate

Not applicable

### Consent for publication

Not applicable

### Availability of data and materials

Sequencing raw reads and processed counts matrices associated with this study will be deposited in the NCBI Gene Expression Omnibus (GEO).

### Authors’ contributions

P.B. and C.J.L conceived the study and wrote the paper. P.B. designed and performed experiments, acquired confocal images, performed data analysis, and generated all figures. P.B. and M.K. designed and performed irradiation experiments and analyses. C.S. performed cloning, riboprobe synthesis for *in-situ* hybridization, and contributed to the intellectual development of the project. N.E.M. conducted statistical analysis for Supplement 1. J.Y. performed the Illumina sequencing. B.W. and C.J.L. contributed to the writing and development of ideas. All authors have read and approved the manuscript.

## Acknowledgements

We would like to thank the staff of the Hopkins Marine Station and the staff of Morro Bay State Park in particular Vince Cicero, John Sayers, and Katie Drexhage for facilitating our collections. We would like to thank David Rank, Paul Peluso, and Greg Conception from Pacific Biosciences, Dan Rokhsar from UC Berkeley, and Norma Neff from Biohub for supporting the development of genomic resources for *Schizocardium*. We thank members of the Lowe Lab, specifically Auston Rutledge who reared many *Schizocardium* larvae and was an invaluable partner in collections. We also thank lab members Laurent Formery, Veronica Pagowski, José Andrade-Lopez, Nat Clarke, Paul Minor, Mark Salvacion, and Catherine Rogers for their helpful discussions. Finally, we particularly thank previous Lowe Lab member Paul Gonzalez for his pioneering work on *Schizocardium*.

## Supplemental Figures

**Supplement 1. Additional characterization of proliferative cells in *S. californicum***

A) Bar chart of number of Hoesch+ in the ciliary bands vs. non ciliary bands, error bars are ±1 SD (66% Confidence interval). B) Bar chart of EdU+ cells in the ciliary bands vs. non ciliary bands, error bars are ±1 SD (66% Confidence interval). C-F) All: anterior up; scale bar is 100um. blue= Hoechst, green= EdU. C) Ventral view of EdU distribution early in metamorphosis. D) Ventral view of EdU distribution in the middle of metamorphosis. E) Lateral view of EdU distribution at the end of metamorphosis. F) Ventral view of EdU distribution in juveniles. G-H) Expression of h2b mRNA and EdU positive cells in the juvenile proboscis blue= Hoechst, yellow= h2b mRNA, green= EdU.

**Supplement 2. Differential expression of larval and juvenile transcriptomes, irradiated versus non-irradiated.**

A) Differential expression of larval transcriptomes with baseMean, log2FoldChange, lfcSE, pvalue, padj. B) Differential expression of juvenile transcriptomes with baseMean, log2FoldChange, lfcSE, pvalue, padj. C) Differential expression of *piwi, vasa,* and *nanos* in larvae with baseMean, log2FoldChange, lfcSE, pvalue, padj. D) Differential expression of *piwi, vasa,* and *nanos* in juveniles with baseMean, log2FoldChange, lfcSE, pvalue, padj.

**Supplement 3. Additional characterization of TUNEL during larval development and metamorphosis**

All: blue = Hoechst, grey= TUNEL, scale bar is 100um. A) lateral of view of mid larval body plan. B) Late larva with very few TUNEL+ cells. C, Highlights regions of B) a few TUNEL+ cells. C) D) Early in metamorphosis from Figure 7C with an increase in TUNEL+ cells. E) TUNEL+ cells found in the mesodermal protocoel. F) Ventral view during the middle of metamorphosis. G) Positive control of TUNEL labeling by artificially nicking the ends of DNA with DNAse-1 metamorphosis H) Inset of positive control with TUNEL detected in deeper tissue layers

**S4. Gene Trees of HCR candidate Genes**

Gene trees for A) Lbr-1 B) Fgfr C) Spne-2

