## Supplement for "Comparisons of cell proliferation and cell death across life histories in the hemichordate *Schizocardium californicum*"

### Supplement 1. Additional characterization of proliferative cells in *S. californicum*

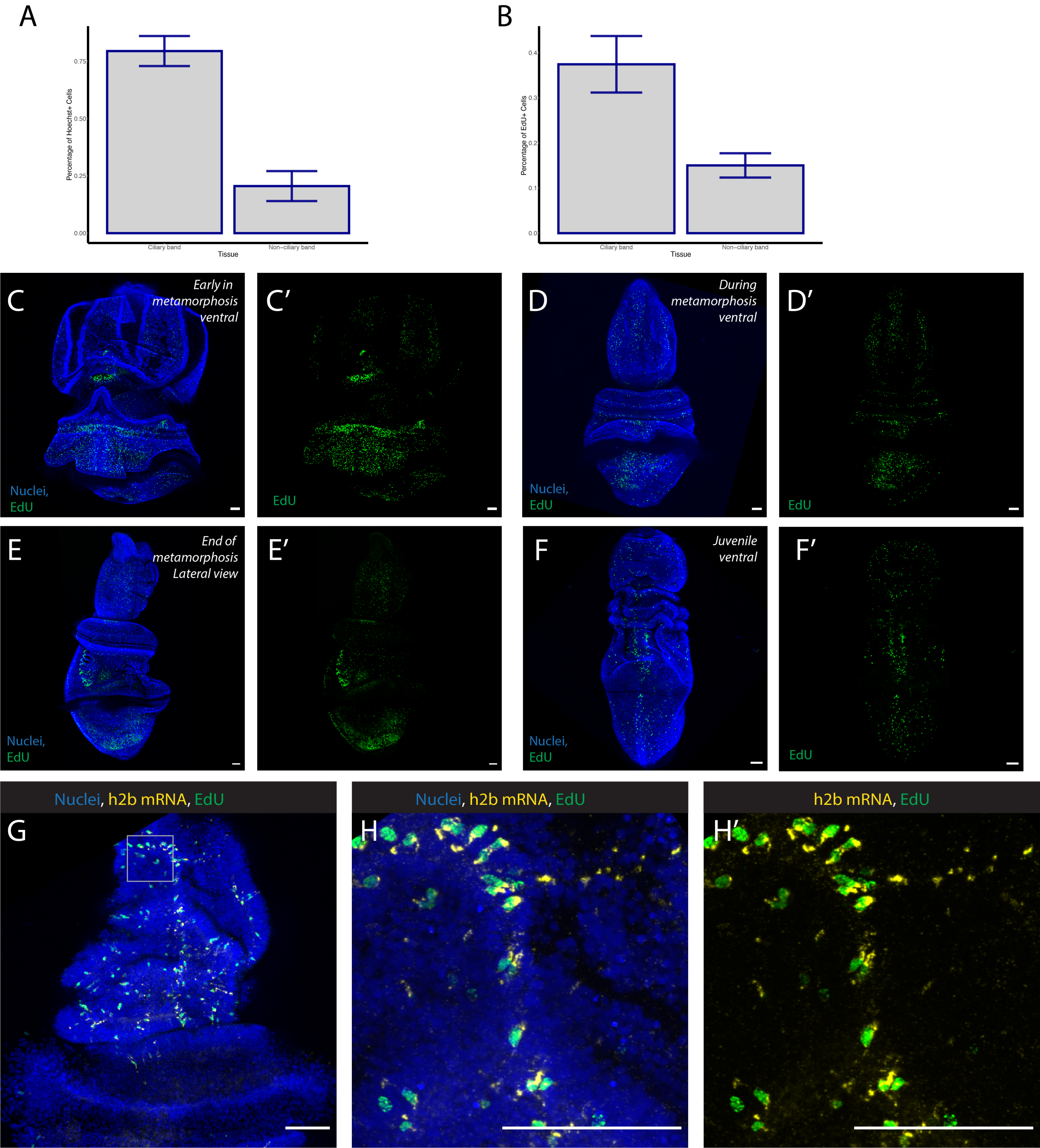

**Supplement 2. Differential expression of larval and juvenile transcriptomes, irradiated versus non-irradiated.**

**A.** Larval Irradiation Candidates

| | baseMean | log2FoldChange | lfcSE | pvalue | padj |
|---|---|---|---|---|---|
| **Fgfr2-4** | 1630.175 | -2.18984 | 0.179364 | 1.04E-35 | 3.34E-32 |
| **Nusap** | 89.84436 | -2.81307 | 0.303536 | 1.09E-21 | 2.33E-18 |
| **Ince** | 119.9263 | -2.22164 | 0.244389 | 5.53E-21 | 8.21E-18 |
| **Aspm-1** | 162.5113 | -2.38698 | 0.268418 | 3.50E-20 | 4.83E-17 |
| **Fgrl1-4** | 91.02211 | -2.31941 | 0.277801 | 3.58E-18 | 3.63E-15 |
| **Mib1-3** | 242.9705 | -2.35456 | 0.288561 | 2.08E-17 | 1.74E-14 |
| **Tlr6-1** | 61.56051 | -2.60842 | 0.323066 | 4.34E-17 | 3.35E-14 |
| **Unchar_3494** | 370.3807 | -2.43616 | 0.310436 | 1.99E-16 | 1.37E-13 |
| **Lbr-1** | 77.78669 | -2.71911 | 0.347314 | 3.07E-16 | 1.98E-13 |
| **Ctk2** | 91.29794 | -2.02464 | 0.288236 | 1.07E-13 | 5.14E-11 |
| **Unchar_1859** | 33.12484 | -2.65533 | 0.378874 | 2.02E-13 | 9.05E-11 |
| **Arrd3-2** | 36.35581 | -3.02327 | 0.446694 | 8.14E-13 | 3.41E-10 |
| **Kif23-1** | 90.78675 | -2.16175 | 0.32543 | 1.76E-12 | 7.21E-10 |
| **Unchar_4293** | 135.4392 | -2.06022 | 0.317639 | 5.08E-12 | 1.92E-09 |
| **Ki11a** | 82.03937 | -2.13209 | 0.336116 | 1.26E-11 | 4.26E-09 |
| **Hmmr** | 41.41535 | -2.26999 | 0.367991 | 3.87E-11 | 1.19E-08 |
| **Dlgp5** | 84.32673 | -2.04988 | 0.336565 | 5.70E-11 | 1.64E-08 |
| **Garem-1** | 50.29227 | -2.20376 | 0.379007 | 4.43E-10 | 1.07E-07 |
| **Tenr-5** | 47.0752 | -2.33731 | 0.442723 | 1.02E-08 | 1.76E-06 |
| **Helz2-7** | 116.1244 | -2.82684 | 0.572553 | 3.50E-08 | 5.19E-06 |

**B.** Juvenile Irradiation Candidates

| | baseMean | log2FoldChange | lfcSE | pvalue | padj |
|---|---|---|---|---|---|
| **Gbp1-2** | 341.796 | -2.4307 | 0.163921 | 6.35E-51 | 2.65E-47 |
| **Ttc22-14** | 145.5675 | -3.19616 | 0.233735 | 1.10E-43 | 3.83E-40 |
| **Neul4-1** | 451.0323 | -2.12873 | 0.159312 | 6.89E-42 | 2.05E-38 |
| **Traf2-30** | 1131.106 | -3.3257 | 0.27781 | 2.92E-34 | 5.08E-31 |
| **Mfha1-3** | 552.3618 | -2.39307 | 0.222611 | 3.92E-28 | 3.71E-25 |
| **Hrsl3-1** | 685.2498 | -2.13015 | 0.199273 | 6.85E-28 | 6.20E-25 |
| **Lecg-2** | 203.9375 | -2.64955 | 0.256702 | 3.95E-26 | 3.04E-23 |
| **Rtbs-11** | 100.8882 | -2.43979 | 0.23934 | 1.28E-25 | 9.17E-23 |
| **Trpa1-12** | 223.4717 | -3.37139 | 0.335112 | 5.49E-25 | 3.47E-22 |
| **Un5cl-2** | 1140.134 | -2.48357 | 0.254042 | 8.08E-24 | 4.43E-21 |

| | | | | | |
|---|---|---|---|---|---|
| **Atp7a-2** | 1473.088 | -2.15204 | 0.224653 | 6.02E-23 | 2.79E-20 |
| **Nckx6-4** | 135.4189 | -3.14621 | 0.337107 | 5.62E-22 | 2.28E-19 |
| **Dzip3-4** | 105.7859 | -2.45534 | 0.2627 | 6.03E-22 | 2.37E-19 |
| **Rtxe-24** | 341.8183 | -3.76393 | 0.411762 | 3.63E-21 | 1.28E-18 |
| **Lrk1-2** | 208.5048 | -3.06119 | 0.341051 | 1.44E-20 | 4.76E-18 |
| **Unchar_3738** | 530.598 | -2.01175 | 0.223771 | 1.49E-20 | 4.85E-18 |
| **Unchar_4293** | 135.4392 | -2.34311 | 0.262811 | 3.71E-20 | 1.17E-17 |
| **Unchar_736** | 239.0824 | -2.02552 | 0.231088 | 1.18E-19 | 3.51E-17 |
| **Tlr6-19** | 75.49353 | -2.78081 | 0.321575 | 4.99E-19 | 1.28E-16 |
| **Unchar_522** | 340.8634 | -2.102 | 0.244208 | 5.14E-19 | 1.31E-16 |
| **Unchar_564** | 772.3732 | -3.5374 | 0.414146 | 6.62E-19 | 1.64E-16 |
| **Par14-36** | 555.8046 | -2.50209 | 0.293003 | 7.23E-19 | 1.77E-16 |
| **Fcgbp-5** | 97.62039 | -2.1658 | 0.257802 | 2.99E-18 | 6.77E-16 |
| **Par14-22** | 499.0506 | -2.64666 | 0.316978 | 3.70E-18 | 8.20E-16 |
| **Gbp3-2** | 51.52854 | -2.62133 | 0.313873 | 6.06E-18 | 1.29E-15 |
| **Neul4-22** | 361.8735 | -2.08839 | 0.251677 | 6.94E-18 | 1.46E-15 |
| **Anln** | 106.3931 | -2.76097 | 0.32943 | 8.41E-18 | 1.72E-15 |
| **Helz2-2** | 471.173 | -3.49158 | 0.425984 | 1.32E-17 | 2.62E-15 |
| **Tlr2-18** | 41.51185 | -3.15919 | 0.38663 | 1.75E-17 | 3.45E-15 |
| **Dtx3l-48** | 690.1412 | -2.1444 | 0.26339 | 2.23E-17 | 4.35E-15 |
| **Tenr-5** | 47.0752 | -2.37845 | 0.293253 | 3.27E-17 | 6.11E-15 |
| **Rn213-22** | 182.4998 | -2.62322 | 0.324544 | 4.26E-17 | 7.72E-15 |
| **Anr55-3** | 100.0866 | -2.62807 | 0.325937 | 4.63E-17 | 8.32E-15 |
| **Sacs-21** | 286.0351 | -2.49152 | 0.314536 | 1.37E-16 | 2.29E-14 |
| **Toll-6** | 295.0914 | -2.03062 | 0.258807 | 3.10E-16 | 4.94E-14 |
| **Znfx1-2** | 920.7174 | -2.33684 | 0.299294 | 3.34E-16 | 5.26E-14 |
| **Ern1-3** | 245.7098 | -2.12359 | 0.272442 | 4.65E-16 | 7.22E-14 |
| **Tsp1-17** | 108.5286 | -3.39356 | 0.436258 | 5.14E-16 | 7.94E-14 |
| **Znfx1-3** | 856.5053 | -2.33028 | 0.301252 | 6.01E-16 | 9.13E-14 |
| **Unchar_3199** | 329.0729 | -2.2087 | 0.288682 | 1.22E-15 | 1.78E-13 |
| **Spne-2** | 87.13935 | -3.61232 | 0.477233 | 2.19E-15 | 3.05E-13 |
| **Nusap** | 89.84436 | -2.57242 | 0.340659 | 3.51E-15 | 4.71E-13 |
| **Tnf10-1** | 199.3844 | -2.3324 | 0.311223 | 4.19E-15 | 5.46E-13 |
| **Gbp1-1** | 57.22204 | -2.65462 | 0.357632 | 6.41E-15 | 8.14E-13 |
| **Ifih1-6** | 239.7474 | -2.87832 | 0.389052 | 7.22E-15 | 9.11E-13 |
| **Par14-28** | 3909.747 | -2.05719 | 0.278852 | 9.81E-15 | 1.19E-12 |
| **Oas3-2** | 51.82069 | -2.61656 | 0.356638 | 1.52E-14 | 1.82E-12 |
| **Otop3-2** | 154.7464 | -2.38236 | 0.326566 | 1.83E-14 | 2.18E-12 |

| | | | | | |
|---|---|---|---|---|---|
| **Helz2-1** | 120.7123 | -2.76886 | 0.379163 | 1.98E-14 | 2.31E-12 |
| **Neul4-30** | 31.62748 | -2.80971 | 0.392604 | 3.92E-14 | 4.41E-12 |
| **Dapk1-19** | 100.0518 | -2.27046 | 0.314664 | 4.09E-14 | 4.56E-12 |
| **Unchar_738** | 493.909 | -2.30478 | 0.327647 | 1.10E-13 | 1.15E-11 |
| **Mx1** | 1858.205 | -2.37222 | 0.33725 | 1.16E-13 | 1.21E-11 |
| **Ddx58-4** | 264.167 | -2.34393 | 0.338489 | 2.61E-13 | 2.57E-11 |
| **Sting-1** | 400.6932 | -2.18297 | 0.316439 | 3.05E-13 | 2.96E-11 |
| **Dtx3l-2** | 42.84706 | -2.16878 | 0.320941 | 8.54E-13 | 7.46E-11 |
| **Sacs-14** | 353.1055 | -2.80702 | 0.41642 | 8.66E-13 | 7.46E-11 |
| **Sacs-15** | 538.5808 | -2.90482 | 0.432026 | 9.06E-13 | 7.74E-11 |
| **Ymd3-16** | 55.01933 | -2.5941 | 0.385597 | 9.79E-13 | 8.26E-11 |
| **Gbp3-1** | 91.71746 | -2.26544 | 0.3364 | 1.01E-12 | 8.45E-11 |
| **Ncan-3** | 123.4335 | -2.02993 | 0.300826 | 1.04E-12 | 8.70E-11 |
| **Arsb-63** | 251.672 | -2.03598 | 0.303042 | 1.12E-12 | 9.29E-11 |
| **Papl-3** | 30.60396 | -3.1363 | 0.469825 | 1.19E-12 | 9.68E-11 |
| **Sacs-9** | 150.0414 | -2.46959 | 0.369306 | 1.38E-12 | 1.10E-10 |
| **Nada-3** | 319.9379 | -2.15271 | 0.32273 | 1.54E-12 | 1.22E-10 |
| **M3k10** | 306.63 | -2.14718 | 0.328022 | 3.46E-12 | 2.56E-10 |
| **Pcsk9-21** | 137.2618 | -2.80255 | 0.439654 | 8.86E-12 | 6.05E-10 |
| **Ncam2-1** | 220.0241 | -2.34591 | 0.371796 | 1.58E-11 | 1.02E-09 |
| **Ddx58-3** | 180.8105 | -2.12169 | 0.339126 | 2.17E-11 | 1.35E-09 |
| **Unchar_922** | 118.2275 | -2.24251 | 0.357785 | 2.55E-11 | 1.57E-09 |
| **Endub-3** | 28.61787 | -2.48477 | 0.399556 | 2.66E-11 | 1.62E-09 |
| **Catl-2** | 181.5082 | -2.74919 | 0.444244 | 3.17E-11 | 1.91E-09 |
| **Ifih1-5** | 252.7791 | -2.02858 | 0.327144 | 3.54E-11 | 2.11E-09 |
| **Arsi-3** | 259.1597 | -2.02039 | 0.328784 | 4.57E-11 | 2.62E-09 |
| **Unchar_3247** | 51.38203 | -2.6797 | 0.431023 | 5.60E-11 | 3.19E-09 |
| **Clca1-6** | 48.81313 | -2.5027 | 0.418174 | 1.16E-10 | 6.14E-09 |
| **Rn213-25** | 21.3758 | -3.01197 | 0.502621 | 1.52E-10 | 7.68E-09 |
| **Unchar_2684** | 171.8979 | -2.12437 | 0.358005 | 1.87E-10 | 9.16E-09 |
| **Trpa1-13** | 37.40464 | -3.0141 | 0.512492 | 2.68E-10 | 1.26E-08 |
| **Par14-35** | 31.14865 | -2.05843 | 0.353369 | 3.26E-10 | 1.48E-08 |
| **Paf15** | 47.28724 | -2.36046 | 0.406903 | 4.59E-10 | 2.02E-08 |
| **Endua-1** | 22.18769 | -2.68102 | 0.464715 | 4.95E-10 | 2.16E-08 |
| **Pgfra** | 21.84278 | -2.82684 | 0.494981 | 5.39E-10 | 2.32E-08 |
| **Helz2-4** | 87.13036 | -2.37221 | 0.414769 | 6.80E-10 | 2.86E-08 |
| **Unchar_2123** | 400.4305 | -2.08215 | 0.372844 | 1.37E-09 | 5.42E-08 |
| **Rn213-23** | 34.64558 | -2.13958 | 0.381366 | 1.51E-09 | 5.88E-08 |

| | | | | | |
|---|---|---|---|---|---|
| **Rab7a-4** | 27.16251 | -2.3785 | 0.427277 | 1.57E-09 | 6.09E-08 |
| **Neul4-38** | 49.2008 | -2.57817 | 0.464363 | 1.76E-09 | 6.74E-08 |
| **Unchar_1989** | 44.5272 | -2.14167 | 0.383324 | 1.77E-09 | 6.75E-08 |
| **Sacs-10** | 66.49401 | -2.18558 | 0.395297 | 2.04E-09 | 7.60E-08 |
| **Herc2-3** | 297.6247 | -2.02755 | 0.368037 | 2.37E-09 | 8.74E-08 |
| **Mb213-5** | 423.577 | -2.13253 | 0.389559 | 2.55E-09 | 9.30E-08 |
| **Cfdp2-64** | 403.5795 | -2.05362 | 0.377091 | 2.74E-09 | 9.97E-08 |
| **Tlr2-16** | 33.61337 | -2.3182 | 0.421844 | 2.88E-09 | 1.04E-07 |
| **Svep1-55** | 39.12176 | -3.55841 | 0.648675 | 3.07E-09 | 1.10E-07 |
| **Lyam3-8** | 20.61618 | -3.52776 | 0.637637 | 3.39E-09 | 1.21E-07 |
| **Pcsk9-28** | 550.1161 | -2.13902 | 0.394645 | 3.39E-09 | 1.21E-07 |
| **Oas3-5** | 30.92615 | -2.90639 | 0.549108 | 4.34E-09 | 1.50E-07 |
| **Lbr-1** | 77.78669 | -2.07277 | 0.38638 | 4.95E-09 | 1.67E-07 |
| **Endua-3** | 34.10096 | -2.88468 | 0.538823 | 6.21E-09 | 2.04E-07 |
| **Tlr2-23** | 38.80898 | -2.50797 | 0.474516 | 8.08E-09 | 2.57E-07 |
| **Arsb-59** | 32.80632 | -2.16404 | 0.412927 | 9.18E-09 | 2.88E-07 |
| **Pcsk9-9** | 72.29667 | -2.11567 | 0.406799 | 1.05E-08 | 3.23E-07 |
| **Tlr2-17** | 18.82426 | -2.36485 | 0.459107 | 1.93E-08 | 5.58E-07 |
| **Trpa1-15** | 32.64893 | -2.96569 | 0.578965 | 2.02E-08 | 5.82E-07 |
| **Tlr6-23** | 19.89627 | -2.32801 | 0.464061 | 3.39E-08 | 9.13E-07 |
| **Tlr6-16** | 31.6292 | -2.40429 | 0.486813 | 4.81E-08 | 1.24E-06 |
| **Helz2-5** | 112.955 | -2.1012 | 0.427976 | 5.25E-08 | 1.34E-06 |
| **Helz2-3** | 775.969 | -2.71311 | 0.559432 | 5.64E-08 | 1.42E-06 |
| **Cfdp2-16** | 21.57929 | -2.41049 | 0.507699 | 9.22E-08 | 2.22E-06 |
| **Susd2-4** | 133.2866 | -2.21769 | 0.466371 | 1.08E-07 | 2.56E-06 |
| **Mb213-7** | 70.49573 | -2.75539 | 0.590883 | 1.22E-07 | 2.87E-06 |
| **Par14-4** | 104.2727 | -2.73857 | 0.588614 | 1.41E-07 | 3.26E-06 |
| **Unchar_2772** | 125.4463 | -2.15169 | 0.463596 | 1.85E-07 | 4.14E-06 |
| **Helz2-8** | 88.94362 | -2.29908 | 0.497874 | 1.95E-07 | 4.33E-06 |
| **Zo1-6** | 39.69926 | -2.39939 | 0.516723 | 2.00E-07 | 4.42E-06 |
| **Anr52-3** | 39.78255 | -2.27627 | 0.493624 | 2.03E-07 | 4.47E-06 |
| **Bgbp-2** | 51.73825 | -2.7608 | 0.600163 | 2.08E-07 | 4.56E-06 |
| **Otop3-1** | 85.54154 | -2.75052 | 0.603507 | 2.45E-07 | 5.26E-06 |
| **Otop2** | 61.2041 | -2.42398 | 0.534917 | 2.84E-07 | 5.98E-06 |
| **Rn213-28** | 805.1838 | -2.56144 | 0.567377 | 2.91E-07 | 6.10E-06 |
| **Cbpa4-1** | 43.58815 | -2.38222 | 0.528439 | 3.05E-07 | 6.33E-06 |
| **Mb213-8** | 45.59932 | -2.91472 | 0.651967 | 3.14E-07 | 6.48E-06 |

**C.** Differential expression of “classic multipotency” genes in larva

| | baseMean | log2FoldChange | lfcSE | stat | pvalue | padj |
|---|---|---|---|---|---|---|
| **Piwi-1** | 93.74719 | 0.39905 | 0.371772 | 1.073373 | 0.283104 | 0.60958 |
| **Nanos** | 998.3336 | -0.60208 | 0.280642 | -2.14537 | 0.031924 | 0.187685 |
| **Piwi-2** | 130.5946 | 0.560572 | 0.277961 | 2.016727 | 0.043724 | 0.226244 |
| **Vasa** | 32.85435 | -0.00313 | 0.434226 | -0.00722 | 0.994243 | 1 |

**D.** Differential expression of “classic multipotency” genes in juveniles

| | baseMean | log2FoldChange | lfcSE | stat | pvalue | padj |
|---|---|---|---|---|---|---|
| **Piwi-1** | 93.74719 | -0.3297 | 0.337528 | -0.9768 | 0.32867 | 0.528603 |
| **Nanos** | 998.3336 | -0.52775 | 0.28077 | -1.87964 | 0.060158 | 0.165342 |
| **Piwi-2** | 130.5946 | 0.463554 | 0.288141 | 1.608776 | 0.107665 | 0.252103 |
| **Vasa** | 32.85435 | -0.113 | 0.459135 | -0.24611 | 0.805593 | 0.89698 |

### Supplement 3. Additional characterization of TUNEL during larval development and metamorphosis

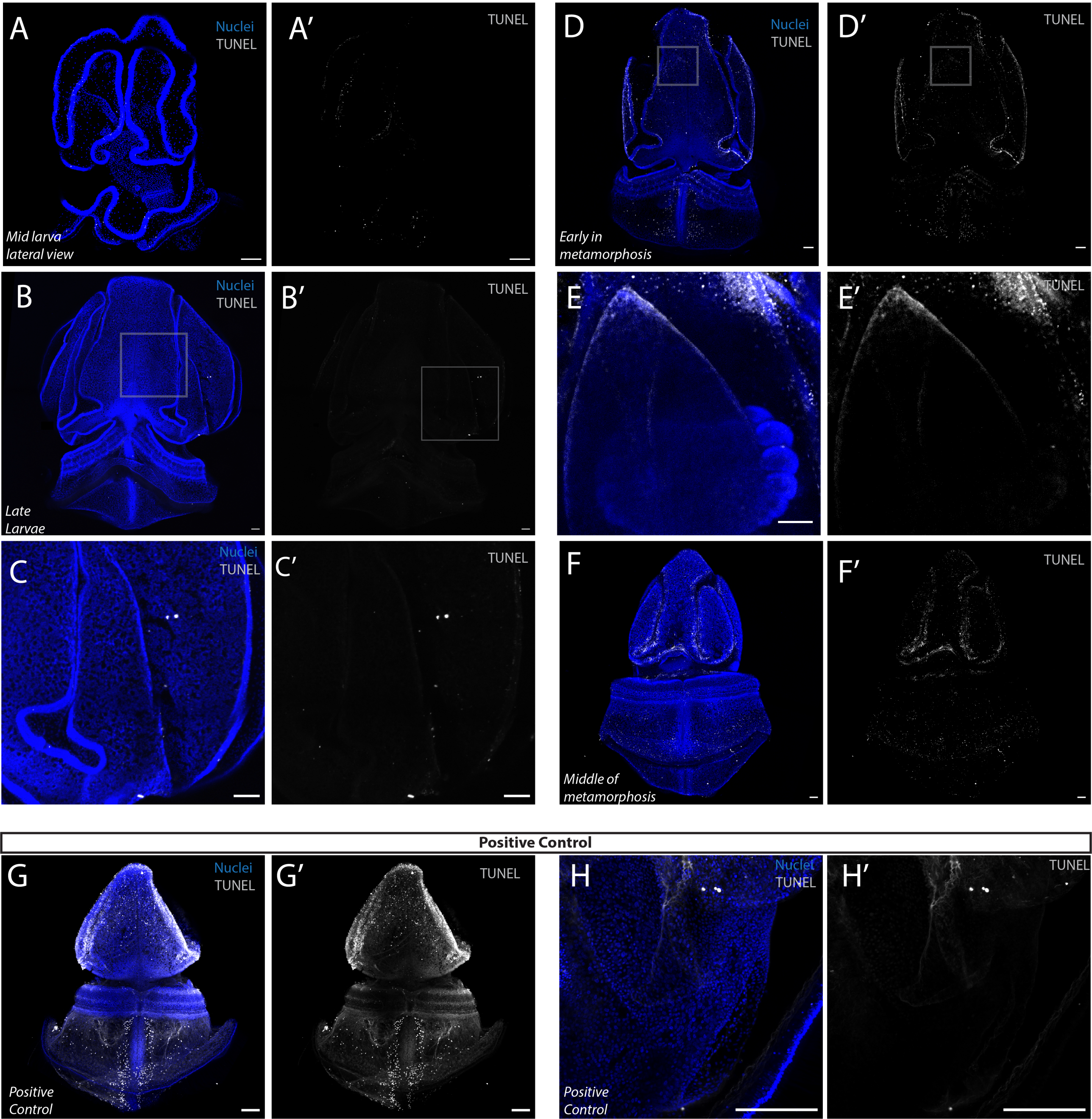

### Supplement 4. Gene Trees of HCR Candidate Genes

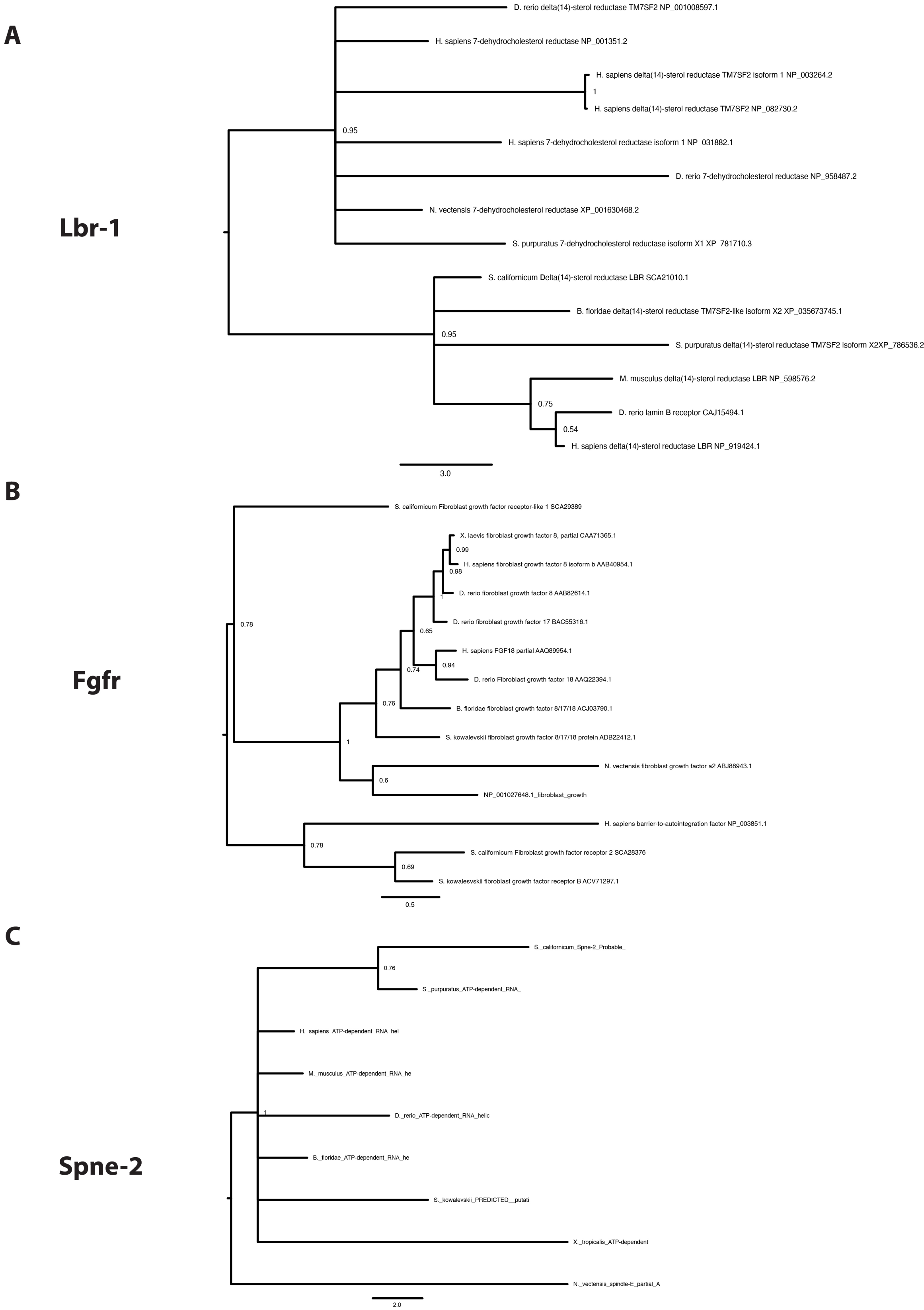
